# *Plasmodium*-encoded murine IL-6 impairs liver stage infection and elicits long-lasting sterilizing immunity

**DOI:** 10.1101/2021.11.16.468835

**Authors:** Selma Belhimeur, Sylvie Briquet, Roger Peronet, Jennifer Pham, Pierre-Henri Commere, Pauline Formaglio, Rogerio Amino, Artur Scherf, Olivier Silvie, Salaheddine Mecheri

## Abstract

*Plasmodium* sporozoites inoculated by *Anopheles* mosquitoes into the skin of the mammalian host migrate to the liver before infecting hepatocytes. Previous work demonstrated that early production of IL-6 in the liver is detrimental for the parasite growth, contributing to the acquisition of a long-lasting immune protection after immunization with live attenuated parasites. Considering that IL-6 ais a critical pro-inflammatory signal, we explored a novel approach whereby the parasite itself encodes for the murine IL-6 gene. We generated transgenic *P. berghei* parasites that express murine IL-6 during liver stage development. Though IL-6 transgenic sporozoites develop into exo-erythrocytic forms in cultured hepatocytes *in vitro* and *in vivo*, these parasites were not capable of inducing a blood stage infection in mice. Furthermore, immunization of mice with transgenic IL-6-expressing *P. berghei* sporozoites elicited a long-lasting CD8^+^ T cell-mediated protective immunity against a subsequent infectious sporozoite challenge. Collectively, this study demonstrates that parasite-encoded IL-6 attenuates parasite virulence with abortive liver stage of *Plasmodium* infection, forming the basis of a novel suicide vaccine strategy to elicit protective antimalarial immunity.

**Summary:** IL-6 was shown to control *Plasmodium* parasite development in the liver. Here, Belhimeur et al. generated a murine IL-6 transgenic *Plasmodium berghei*. These parasites show an arrest in hepatocyte development and protect mice against homologous and heterologous parasite challenge in a CD8-dependent manner.

## Introduction

Major efforts have been made to design vaccines against malaria, an infectious disease that causes serious economic and health problems [1]. With increasing emergence worldwide of resistant *Plasmodium* parasite strains to antimalarial drugs, vaccines remain a critical strategy to control malaria. In recent years, in addition to the development of subunit vaccines and radiation-attenuated sporozoites (SPZ) (RAS), researchers have used rodent models to test the efficacy of genetically attenuated parasites (GAPs) as vaccines against exo-erythrocytic forms (EEF) and blood-stage malaria infections. Most studies have focused on EEF of GAPs, which include parasite mutants blocked early during their development in the liver. Among them, parasites lacking integrity of the parasitophorous vacuole, and parasites deficient for fatty acid biosynthesis type II pathway [2] have been reported. A new concept has emerged from these studies that replication competent and late stage-arresting GAPs were more efficient in generating sterile protection against malaria when compared with replication-deficient and early liver stage-arresting GAPs [1].

Few molecules of malaria parasites have been shown to counteract host innate immunity. In early liver stages, the major SPZ surface protein called circumsporozoite protein (CSP) translocates into the hepatocyte cytosol and nucleus, where it outcompetes NF-kB nuclear import and suppresses hundreds of genes involved in the host inflammatory response [3]. It has also been reported that several parasites including *Plasmodium*, *Brugia*, *Trichuris*, *Eimeiria*, *Trichinella*, and *Onchorcerca*, express a *migration inhibitory factor* (MIF) orthologue of mammalian *MIF* with significant homology to human *MIF*. MIF was shown to increase inflammatory cytokine production during the blood phase of *Plasmodium* infection in rodents and to induce antigen-specific CD4 T-cells to develop into short-lived effector cells rather than into memory cells, causing decreased CD4 T-cell recall responses to homologous parasites [4]. However, parasites lacking MIF were shown to have no growth defect throughoutthe parasite life cycle in *P. berghei* (*Pb*) [5] or a growth defect during liver-stage development in *P. yoelii* [6]. We have investigated the protection mediated by an erythrocytic GAP depleted of the gene encoding the immunomodulatory and secreted molecule histamine releasing factor (HRF), using the parasite strain *Pb*NK65 that does not cause experimental cerebral malaria (ECM) and rapid death. In earlier work, we found that blood-stage infection by the mutant self-resolved at day 12 post-infection (p.i.), displaying an immune signature that comprised elevated IL-6 levels, activation of T and B cells, and antigen-specific IgG2c production [7]. During the last decade, in search of key mechanisms that determine the host inflammatory response, a set of host factors turned out to be critical for the control of malaria parasite development [8, 9, 7, 10]. Strikingly, the activation of the IL-6 signaling pathway occurred as a common theme, that apparently blocks parasite liver stage development, but more importantly allows the initiation of a robust and long-lasting anti-parasite immunity.

In this report we explored a novel approach whereby the parasite itself encodes for the murine *IL-6* gene. This strategy is based on the rational that IL-6 acts as a critical pro-inflammatory signal from the host in response to the parasite infection. We developed a novel suicide strategy to restrict *Plasmodium* liver stage development through parasite-encoded murine *IL6*. We engineered transgenic *Pb* parasites to express and secrete high levels of the murine IL-6 cytokine restricted to the liver stage and evaluated their potential to cause an attenuated infection and to confer protection against wild-type virulent parasite challenges. We demonstrate that parasite-encoded *IL-6* efficiently attenuates *Plasmodium* infectivity at the liver stage and elicits a long-lasting cross-species protective antimalarial immunity.

## Results

### Transgenic *P. berghei* parasites expressing IL-6 during the liver stage lose infectivity to mice

To generate IL-6 expressing transgenic parasites, we integrated an *IL-6* cassette by homologous recombination in the genome of *P. berghei* ANKA lines, which constitutively express the green fluorescent protein (GFP) [11] (Fig. 1A). For this purpose, a codon-optimized *IL-6* gene comprising amino acids 25-211 of murine IL-6 was fused to the signal peptide of *P. berghei* liver-specific protein 2 (LISP2), to enable secretion of the protein. The resulting transgene was placed under the control of *LISP2* promoter to ensure timely restricted expression to the liver stage development [12, 13]. To facilitate integration of the construct, we used a selection-linked integration strategy [14] to insert the *IL-6* cassette into the GFP locus of *Pb*GFP parasites (Fig. 1A). Following transfection, one single round of pyrimethamine selection was sufficient to recover pure populations of transgenic parasites. Two transfectants were selected from independent transfection experiments, called IL-6 Tg-*Pb*ANKA/LISP2 line 1 and line 2, and were confirmed by PCR using the primer combinations shown in Suppl. Table 1, to harbor the expected integration of the murine *IL-6* gene in a similar way (Fig. 1B).

**Figure 1:**
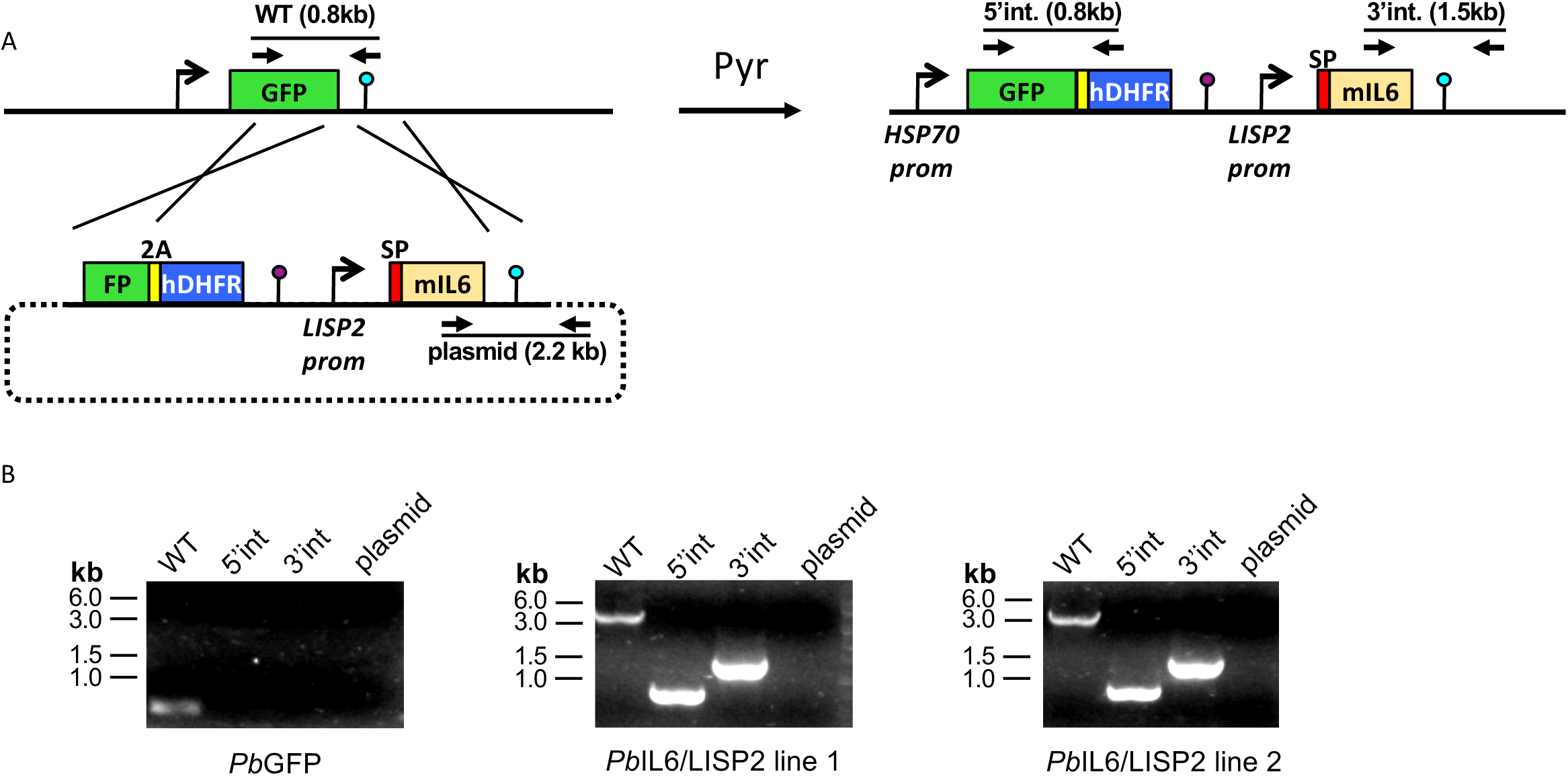
Design of transgenic *Plasmodium berghei* parasites that express the mouse IL-6 cytokine. A) Replacement strategy to modify the *Pb*GFP locus to express murine IL6. We used a selection-linked integration strategy to integrate a construct containing a GFP-2A-hDHFR cassette without promoter followed by the mIL6 cassette under control of the *PbLISP2* promoter. The signal peptide (SP, red) of *Pb*LISP2 was used to allow IL6 secretion. Parasites undergoing homologous recombination at the GFP and *Pb*DHFR 3’UTR sequences were isolated after one round of pyrimethamine selection. Genotyping primers and expected PCR fragments are indicated by arrows and lines, respectively. B) PCR analysis of genomic DNA from parental *Pb*GFP, *Pb*IL6/LISP2 line 1 and line 2 parasites. We used primer combinations to detect the non-recombined parental locus (WT, 830-bp), the 5’ integration event (5’int, 819-bp), the 3’ integration event (3’int, 1554-bp) or the non-integrated construct (plasmid, 2197-bp). The WT primer combination amplifies a 4259-bp fragment from the recombined locus of *Pb*IL6/LISP2 parasites. The results confirm the correct integration of the constructs at the GFP locus of *Pb*GFP parasites, and show the absence of parental parasites in the transfectants after a single round of pyrimethamine selection.

**Supplementary Table 1:**
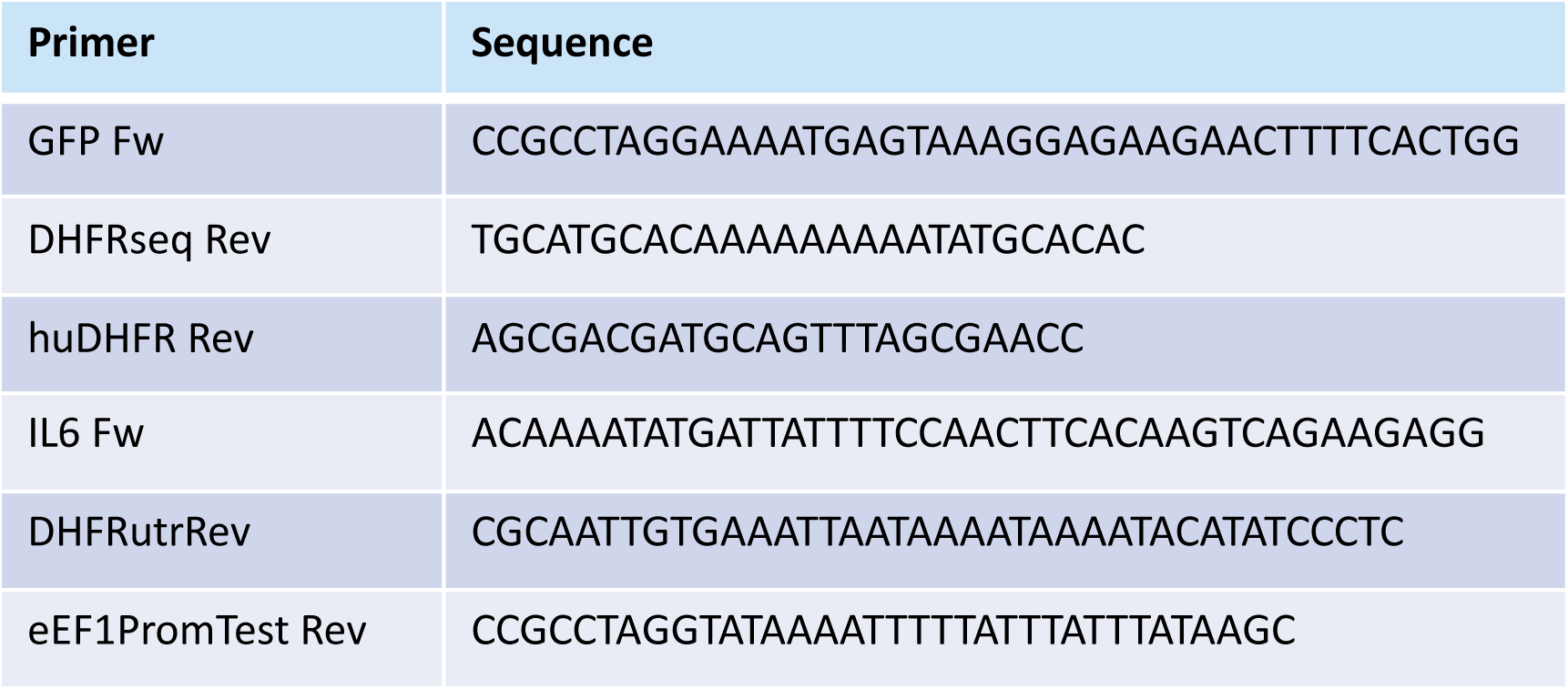
primer combinations to assess the integration of the murine IL-6 transgene into the *Pb*ANKA parasite genome.

When IL-6 Tg-*Pb*ANKA/LISP2 and WT *Pb*ANKA parasites were transmitted to female *Anopheles* mosquitoes, we observed similar prevalence of mosquito infection (Fig. 2A) and numbers of salivary gland SPZ (Fig. 2B). This indicates that the presence of the *IL-6* transgene does not alter parasite development inside mosquitoes. The two lines behave similarly.

**Figure 2:**
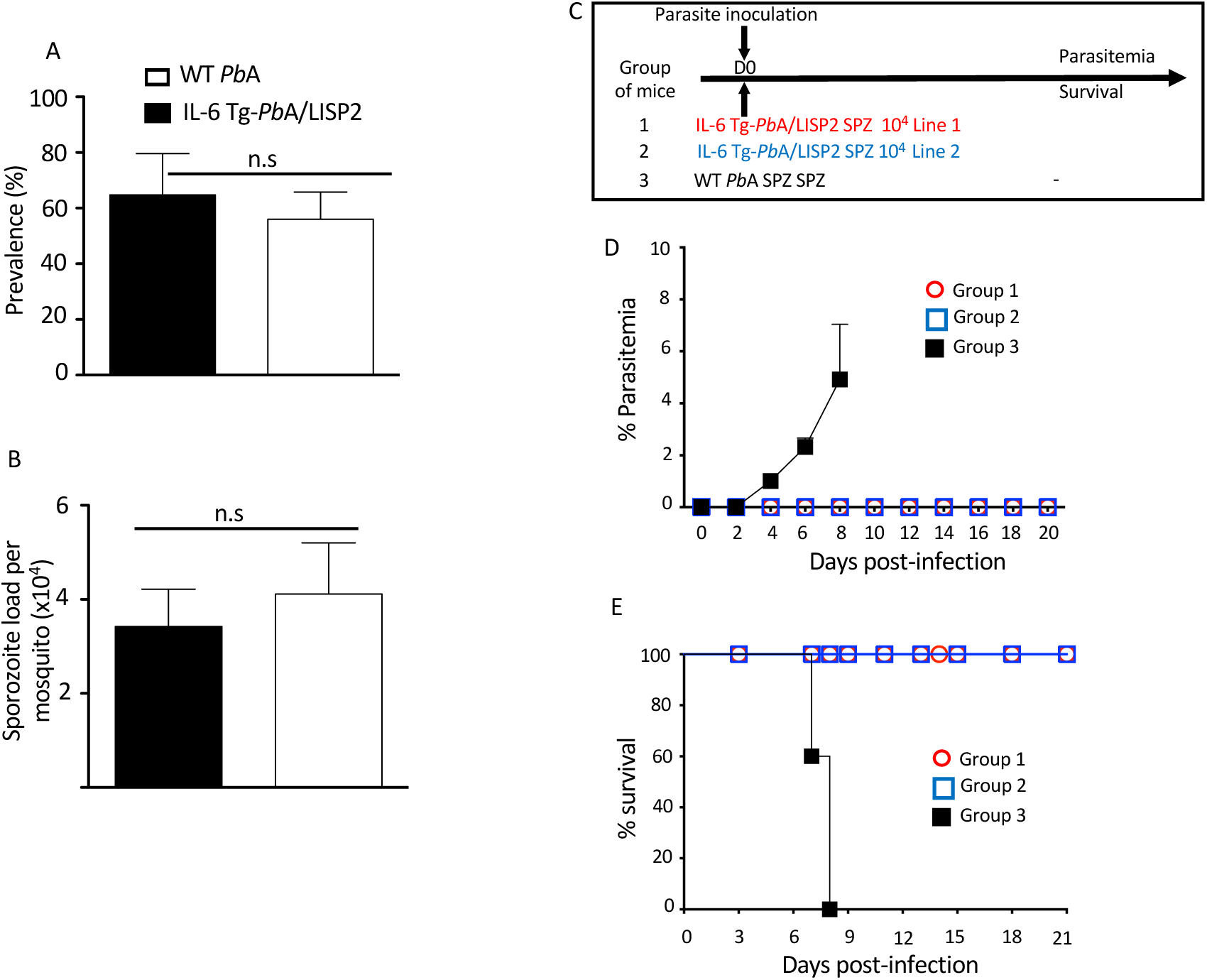
IL-6 Tg-*Pb*ANKA/LISP2 did not show any major development defect in the mosquito vector whereas inoculation with IL-6 Tg-*Pb*A/LISP2 SPZ does not produce blood stage infections. A, B) IL-6 Tg-*Pb*A/LISP2 parasites cycle normally between the vertebrate host and *Anopheles* mosquitoes. Cages of 200 IL-6 Tg-*Pb*A/LISP2- or WT *Pb*ANKA-infected *Anopheles stephensi* female mosquitoes showing GFP-labelled SPZ in their salivary glands were counted at day 20 post-blood feeding on infected C57BL/6 mice. Prevalence of infected mosquitoes was expressed as a ratio between positive ones and total mosquitoes (A). SPZ were extracted from salivary glands of 10 mosquitoes and counted. The number of SPZ was expressed as per pair of salivary glands (B). C) Three groups of C57BL/ mice were infected with 10^4^ IL-6 Tg-*Pb*A/LISP2 SPZ line 1 or with IL-6 Tg-*Pb*A/LISP2 SPZ line 2 or with WT *Pb*ANKA SPZ. D) Parasitemia were measured at indicated time points by flow cytometry, as all parasites were GFP tagged. E) Survival rates were determined by Kaplan-Meier survival plots. Error bars, SEM. Data are representative of three independent experiments with 5 mice per group.

To examine the capacity of IL-6 transgenic sporozoites (SPZ) to develop *in vivo* inside hepatocytes and to differentiate into blood stages, C57BL/6 mice were injected i.v. with 10^4^ SPZ of IL-6 Tg-*Pb*ANKA/LISP2 line 1, or line 2, or with parental WT (*Pb*GFP) SPZ, as shown in Fig. 2C, and parasitemia was followed over time. Parasitemia was determined by flow cytometry by counting GFP^+^ infected red blood cells. As shown in Fig. 2D, we could observe that while mice infected with WT *Pb*ANKA parasites all developed parasitemia, mice that were inoculated with IL-6 Tg-*Pb*ANKA/LISP2 line 1 and line 2 SPZ all failed to show any detectable parasitemia over a period of 20 days of follow up. Mice infected with WT *Pb*ANKA parasites developed experimental cerebral malaria (ECM) and died around day 8 post-infection whereas mice injected with either IL-6 Tg-*Pb*ANKA/LISP2 line 1 or line 2 survived (Fig. 2E). These data demonstrate that insertion of the mouse *IL-6* gene in *Pb*ANKA parasites under control of a liver stage specific promoter results in the complete blockade of parasite development following SPZ injection. From now on, only the IL-6 Tg-*Pb*ANKA/LISP2 line 1 was used throughout the study.

### IL-6 Tg-*Pb*ANKA/LISP2 parasites develop in cultured hepatocytes but arrest during liver stage development in mice

Next, mice were injected with SPZ i.v. and liver samples collected at indicated time points were subjected to RT-qPCR analysis of parasite 18S rRNA (Fig. 3A). The WT *Pb*ANKA parasite liver loads increased over time to reach a maximum at 48 h, followed by a decrease at 72 h, reflecting complete liver stage development and egress. In contrast, the liver load of IL-6 Tg-*Pb*ANKA/LISP2 parasites remained low over the course of infection, except at the 48-h time point, when the amount of IL-6 Tg-*Pb*ANKA/LISP2 parasites rose but at levels significantly lower than those obtained for WT *Pb*ANKA parasites. Therefore, RT-qPCR analysis in the liver confirms that the loss of infectivity of IL-6 Tg-*Pb*ANKA/LISP2 SPZ is due to a defect in liver stage development *in vivo*.

**Figure 3:**
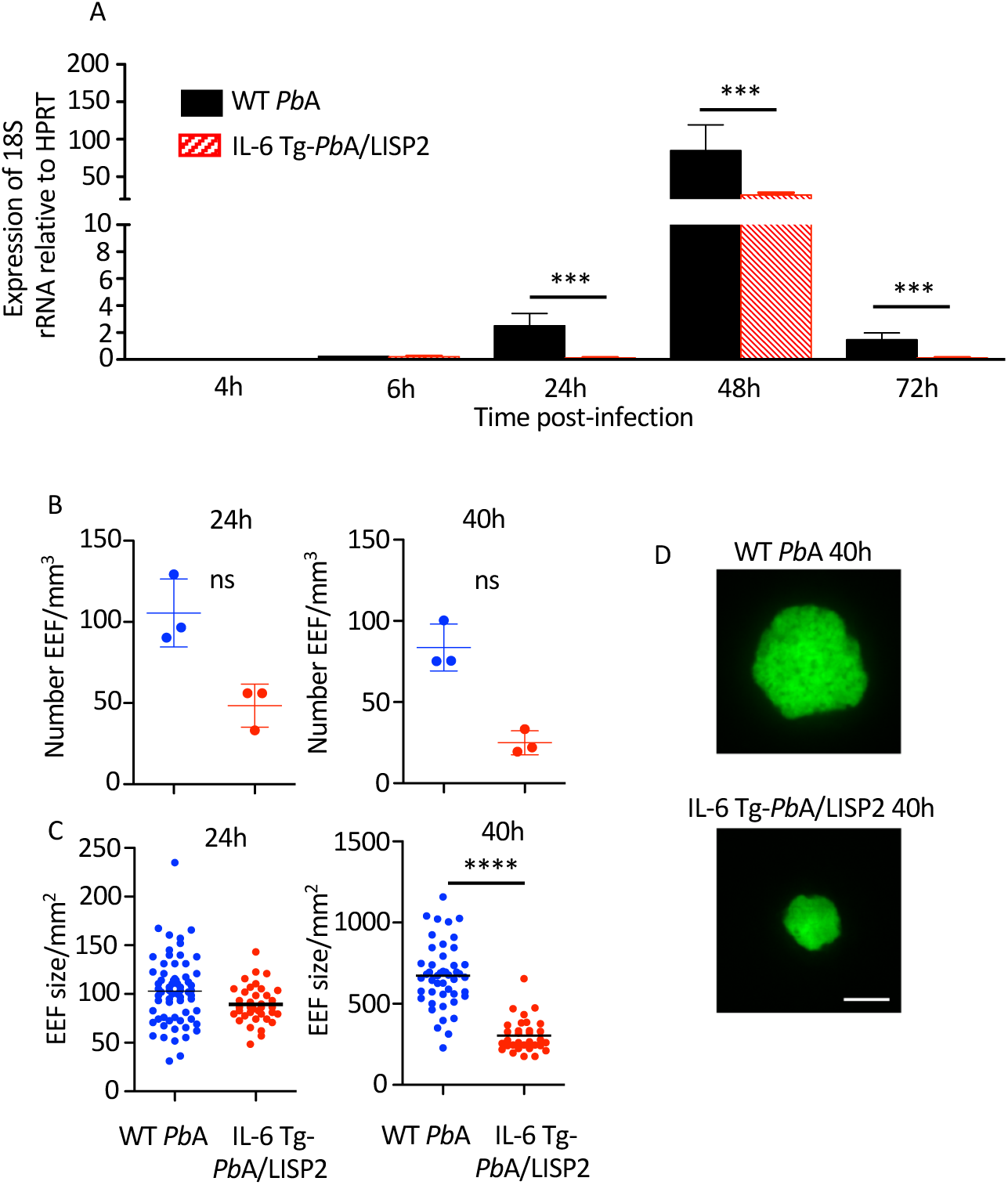
IL-6 Tg-*Pb*ANKA/LISP2 did not show any major development defect in cultured hepatocytes, in contrast to a deficient growth in the liver of infected mice. A) Groups of C57BL/6 mice were injected i.v with 10^4^ WT *Pb*ANKA or with IL-6 Tg-*Pb*ANKA/LISP2 SPZ and livers were collected at indicated time points. Parasite loads in the liver were assessed by measuring parasite 18S rRNA by real-time RT-qPCR. Gene mRNA expression was normalized to the parasite control gene HPRT Data are presented as the means ± SD from six individual values (***P < 0.001; Mann Whitney test). (B-D) Groups of 3 C57BL/6 mice were injected i.v with 150,000 WT *Pb*ANKA or IL-6 Tg-PbANKA/LISP2 SPZ and livers were harvested and imaged at the indicated time points. B) EEF density was determined for each condition and time point analyzing 8-25 Z-stacks per liver using a 10x objective. Bars denote the mean and standard deviation (Kruskall-Wallis test followed by Dunn’s multiple comparison, ns: non significant). EEF area (C) was determined on maximal intensity projections of the acquired Z-stacks using Fiji. Graphs show pooled data from 3 livers per condition and time point (n= 32-66 per group), horizontal bars denote the mean (One-way ANOVA followed by Bonferroni post-test, **** p<0.0001, ns: non significant). D) Representative images of WT *Pb*ANKA or IL-6 Tg-PbANKA/LISP2 EEF at 40 hours post-infection. Scale bar: 10 µm.

To confirm that IL6 expression impairs liver stage development of transgenic parasites, we examined by fluorescence microscopy exoerythrocytic forms (EEFs) in the liver of mice harvested 24h or 40h after infection with 5 x 10^4^ WT *Pb*A or IL-6 Tg-*Pb*ANKA/LISP2 SPZ. As shown in Fig. 3B, the EEF density was drastically reduced both at 24h and at 48h in mice infected with the IL-6 transgenic parasites as compared to those infected with WT *Pb*A parasites, although the differences were not statistically significant. We also examined the size (Fig. 3C) of EEFs, and found the same tendency, namely a reduced size and diameter of IL-6 transgenic EEF as compared to those of WT *Pb*A EEFs with a statistical difference only at 40h. Representative images of WT *Pb*ANKA or IL-6 Tg-*Pb*ANKA/LISP2 EEF at 40 hours post-infection are shown in Fig. 3D. Altogether, these results suggest that IL-6 transgenic parasites display a developmental defect at the pre-erythrocytic stage *in vivo*.

Considering the differential intrahepatic development of WT and IL-6 recombinant parasites, concurrent mouse infection with the two parasites was performed to examine how they interfere with each other’s development. Mice were infected with a mixture of 10, 000 GFP-tagged IL-6 Tg-*Pb*ANKA/LISP2 and mCherry-tagged WT *Pb*A SPZ, and parasitemia and survival were monitored over time. As expected, mice which received either GFP-tagged IL-6 Tg-*Pb*ANKA/LISP2 (Suppl. Fig. 1, filled green circles) or mCherry-WT *Pb*A SPZ (Suppl. Fig. 1, filled red squares) failed to develop parasitemia or showed a patent parasitemia, respectively. Surprisingly, mice co-inoculated with WT and recombinant parasites showed that both red fluorescent WT *Pb*A (filled black squares) and green fluorescent IL-6 Tg-*Pb*ANKA/LISP2 (open green circles) developed into blood stage parasites (Suppl. Fig. 1). These data demonstrate that co-infection with WT parasites reversed the infection phenotype of the IL-6 recombinant parasite, and suggest that the anti-IL-6 signaling provided by WT parasites is dominant over the IL-6 signaling expected to result from IL-6 transgenic parasites.

### IL-6 Tg-*Pb*ANKA/LISP2 parasites express *IL-6* mRNA and secrete the murine IL-6 cytokine

We next assessed whether transgenic parasites express IL-6 in the liver of infected mice. The livers of C57BL/6 mice inoculated with 10,000 SPZ of either IL-6 Tg-*Pb*ANKA/LISP2 or WT *Pb*ANKA parasites were harvested at 48h post-infection and were subjected to RT-qPCR using primers designed to detect the codon-optimized parasite-encoded IL-6 transgene. As shown in Fig. 4A, *IL-6* transgene mRNA could be detected in mice infected with IL-6 Tg-*Pb*ANKA/LISP2 parasites. To verify whether the IL-6 Tg-*Pb*ANKA/LISP2 parasites secrete the IL-6 cytokine, supernatants were collected 48h after the HepG2 hepatoma cell line was cultured in the presence of IL-6 Tg-*Pb*ANKA/LISP2, or with WT *Pb*ANKA SPZ. IL-6 measured by ELISA was present only in supernatants from IL-6 Tg-*Pb*ANKA/LISP2 / HepG2 cultures, but not from WT *Pb*ANKA / HepG2 cultures (Fig. 4B). These data demonstrate that IL-6 transgenic parasites are able not only to express *IL-6 in vitro* and *in vivo*, but also to secrete IL-6, and suggest that the IL-6 transgene is fully functional, likely restricting liver stage development *in vivo*.

**Figure 4:**
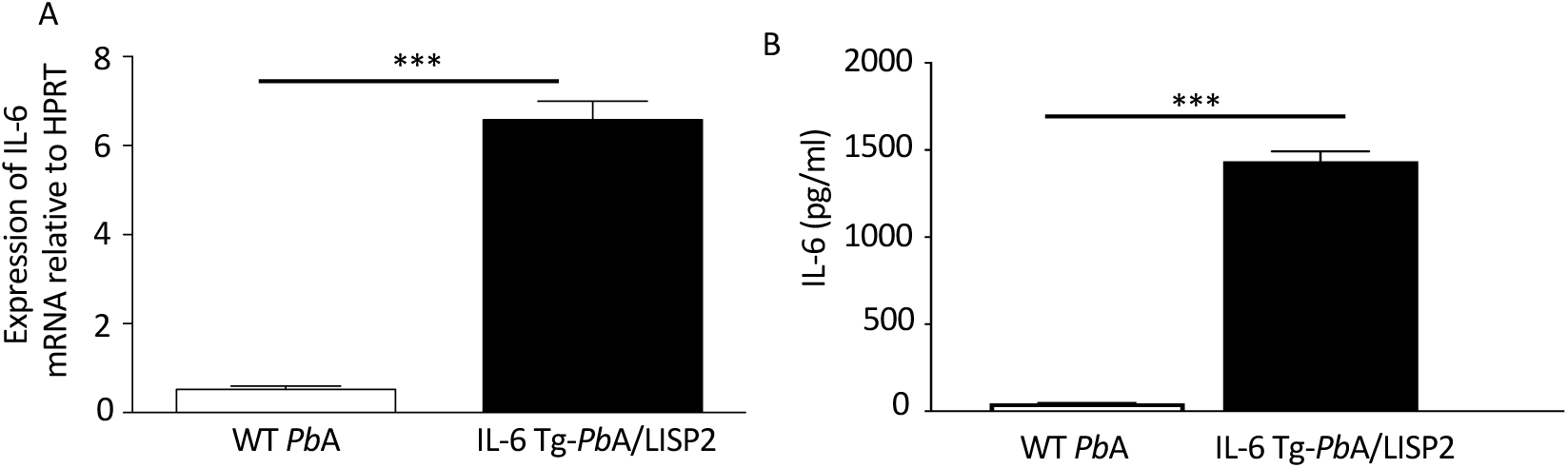
IL-6 Tg-*Pb*ANKA/LISP2 parasites express IL-6 mRNA and produce the murine IL-6 cytokine. A) After infection with WT *Pb*ANKA or IL-6 Tg-*Pb*ANKA/LISP2 SPZ, livers were collected at 48h post-infection, and RT-qPCR analysis was used to measure the IL-6 mRNA expressed by EEFs using a dedicated set of primers relative to the parasite control gene HSP70. B) Determination of IL-6 by ELISA in supernatants obtained from HepG2 cultured in the presence of IL-6 Tg-*Pb*ANKA/LISP2 parasites or WT *Pb*ANKA for 48h. IL-6 was found to be secreted only in cultures with IL-6 Tg-*Pb*ANKA/LISP2 SPZ. Error bars, SEM. Data are representative of two independent experiments using triplicate samples (***p < 0.001; Mann Whitney test).

Interestingly, treatment of mice with anti-IL-6 receptor blocking antibodies before, during and after inoculation of IL-6 Tg-*Pb*ANKA/LISP2 SPZ (Suppl. Fig. 2A), was not sufficient to reverse the developmental defect of transgenic parasites neither in terms of parasitemia (Suppl. Fig. 2B) nor survival (Suppl. Fig. 2C), suggesting that IL-6 produced by transgenic parasites very likely exerts its signaling effects intracellularly in infected hepatocytes.

### Infection with IL-6 Tg-*Pb*ANKA/LISP2 parasites confers protection against challenge with WT *Pb*ANKA sporozoites

We decided to investigate whether the abortive development in mice of IL-6 Tg-*Pb*ANKA/LISP2 parasites could result in protection against secondary infection with WT *Pb*ANKA SPZ. For this, different groups of mice were infected with IL-6 Tg-*Pb*ANKA/LISP2 SPZ using either 10^4^ SPZ (group 1) or 5 x 10^4^ SPZ (group 2) followed by a challenge with 10^4^ WT *Pb*ANKA SPZ at day 30 post-infection with IL-6 Tg parasites (Figure 5A). Control groups consisted in mice infected with 10^4^ or 5 x 10^4^ WT *Pb*ANKA SPZ at day 0 (groups 3 and 4) and mice that received 10^4^ WT *Pb*ANKA SPZ at day 30 (group 5). As expected, all control mice that received only WT *Pb*ANKA SPZ, either at day 0 (Groups 3 and 4, Fig. 5B, C) or at day 30 (group 5) developed parasitemia and died. In contrast, mice previously infected with IL-6 Tg-*Pb*ANKA/LISP2 parasites displayed low parasitemia at day 6 post-challenge, irrespective of the dose of parasites (Groups 1 and 2), with a delay in the pre-patent period of 2 days (day 36 versus day 34, highlighted by the log scale representation of the data) as compared to control mice infected with WT *Pb*ANKA SPZ (Group 5). This delay represents approximately 2 Log difference (100-fold) in parasite load between control and protected mice. Consistently, no mice which received IL-6 Tg-*Pb*ANKA/LISP2 SPZ died from ECM (Fig. 5C). Similar observations were made when secondary infection with WT *Pb*ANKA SPZ was done at 60 days post priming (Suppl. Fig. 3A). IL-6 Tg-*Pb*ANKA/LISP2 SPZ-immunized mice (Groups 1 and 2) were protected from ECM but however later succumbed from hyperparasitemia (Suppl. Fig. 3B, C). These data demonstrate that one single inoculation with IL-6 transgenic *Pb*ANKA/LISP2 parasites is efficient in protecting mice against disease, but is not sufficient to induce sterile protection, even at the high dose of 5 x 10^4^ SPZ.

**Figure 5:**
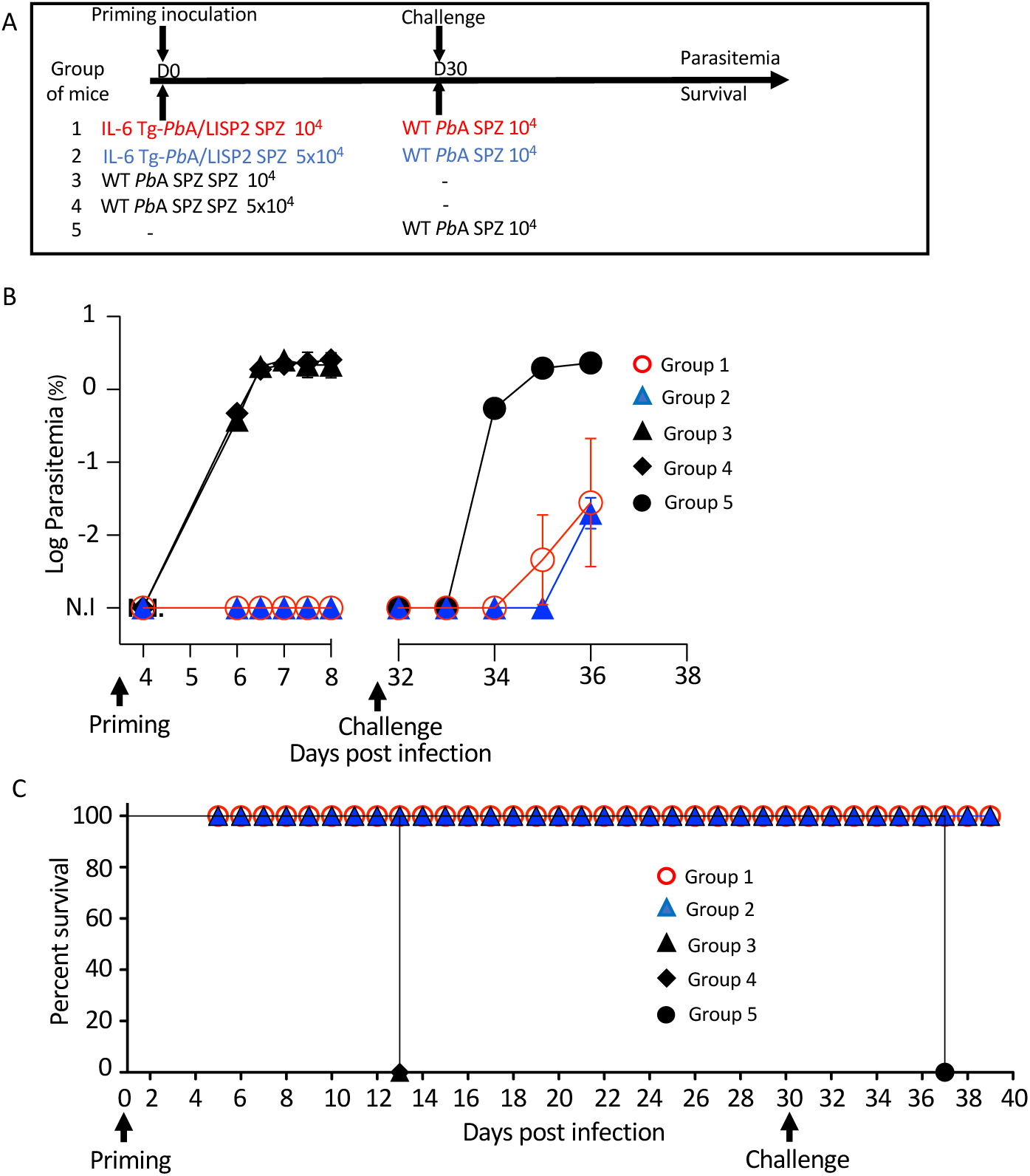
a single inoculation with IL-6 Tg-*Pb*A/LISP2 SPZ is partially efficient in protecting mice against a lethal challenge. A) Groups of C57BL/6 mice were infected with either 10^4^ or 5 x 10^4^ IL-6 Tg-*Pb*A/LISP2 SPZ or with the same doses of WT *Pb*ANKA SPZ. Mice were then challenged with 10^4^ *Pb*ANKA SPZ 30 days after priming. B) Parasite development was measured at indicated time points by flow cytometry, as all parasites were tagged with GFP. C) Survival rates were determined by Kaplan-Meier survival plots. Error bars, SEM. Data are representative of three independent experiments with 5 mice per group.

To optimize the protocol of the IL-6 Tg parasite-mediated protection, additional experiments were conducted in which mice were infected twice with 10^4^ IL-6 Tg-*Pb*ANKA/LISP2 SPZ at three weeks interval, and then challenged 30 days (Fig. 6A) or 60 days (Suppl. Fig. 3A) later with 10^4^ WT *Pb*ANKA SPZ. As shown in Figure 6B, mice that were infected with 10^4^ IL-6 Tg-*Pb*ANKA/LISP2 SPZ (group 1) not only did not show any parasitemia after the first and the second infection with the recombinant parasites, but more interestingly did not develop any parasitemia upon later infection with WT parasites. Using the same infection scheme, we examined whether the protection against a lethal challenge depends on the dose of the primary IL-6 Tg-*Pb*ANKA/LISP2 infection. Groups of mice were infected twice at three weeks interval with IL-6 Tg-*Pb*ANKA/LISP2 SPZ at 10^3^, or 10^2^ SPZ per mouse followed 30 days later by a challenge with 10^4^ WT *Pb*ANKA SPZ (Fig. 6A). As shown in Fig. 6B, the two doses of 10^3^, or 10^2^ of IL-6 Tg-*Pb*ANKA/LISP2 SPZ (group 2 and group 3, respectively) were inefficient at inducing sterile protection as mice developed parasitemia at the same magnitude as the control mice, which received WT *Pb*ANKA SPZ only. All mice that had received the doses of 10^4^ and 10^3^ of IL-6 Tg-*Pb*ANKA/LISP2 SPZ, and 40% of mice injected with the dose of 10^2^ SPZ, were protected from ECM (Fig. 6C). In a long-term follow up, all mice immunized twice with 10^4^ IL-6 Tg-*Pb*ANKA/LISP2 SPZ resulted in sterile protection and survived, while those treated with the 10^3^ and 10^2^ doses of IL-6 Tg-*Pb*ANKA/LISP2 SPZ ultimately died from hyperparasitemia at day 9 and 10 after challenge, respectively. These data indicate that the prime boost immunization at the dose of 10^4^ IL-6 Tg-*Pb*ANKA/LISP2 SPZ was optimal in conferring both clinical and sterilizing immunity.

**Figure 6:**
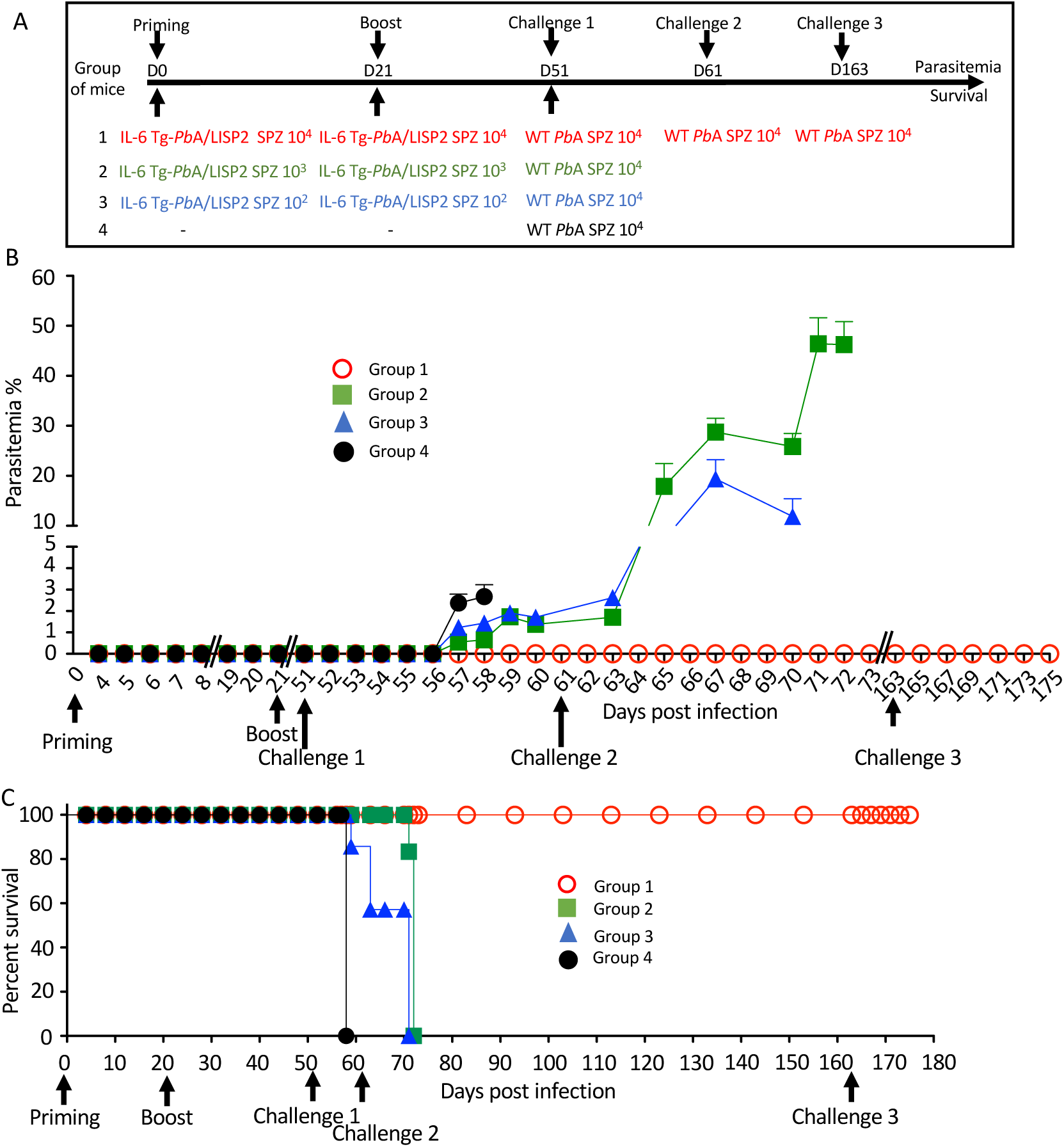
homologous prime/boost immunization regimen with IL-6 Tg-*Pb*ANKA/LISP2 parasites confers a stable and efficient protection against challenge with WT *Pb*ANKA parasites. A) Groups of C57BL/6 mice were infected with various doses of IL-6 Tg-*Pb*A/LISP2 SPZ and boosted with the same doses at 3 weeks interval. Mice were then challenged 30 days after the boost (day 51) with 10^4^ WT *Pb*ANKA SPZ. Mice of group 1, which all survived the lethal challenge, were then re-challenged at day 61 and day 163 with 10^4^ *Pb*ANKA SPZ. B) Parasite development was measured at indicated time points by flow cytometry, as all parasites were tagged with GFP. C) Survival rates were determined by Kaplan-Meier survival plots. Error bars, SEM. Data are representative of three independent experiments with 5 mice per group.

In order to explore how long this protection could last, a slightly different protocol (Suppl. Fig. 4A) was carried out where mice which received the priming and booster injections with IL-6 Tg-*Pb*ANKA/LISP2 SPZ were challenged 60 days after the boost. Similar results were obtained in terms of parasitemia (Group 1, Suppl. Fig. 4B) and survival (Group 1, Suppl. Fig 4C) as in the mice challenged 30 days after the second IL-6 Tg-*Pb*ANKA/LISP2 infection, shown in Fig. 6, with a sterile protection against a challenge with WT *Pb*ANKA SPZ in a homologous prime/boost immunization regimen (Group 1, Suppl. Fig. 4A). This sterile protection which occurred 30 or 60 days after the second IL-6 Tg-*Pb*ANKA/LISP2 SPZ infection strongly suggests the induction of a robust memory response. To assess how efficient was the anti-malaria protection induced by IL-6 transgenic parasites, protected mice already challenged with WT *Pb*ANKA at day 51, were re-challenged twice with WT *Pb*ANKA at day 61 and later at day 163 (Group 1, Fig. 6A). As shown in Fig. 6 B, and C, none of the mice of this group developed parasitemia and all mice survived, indicating that protected mice survived and remained parasite free after multiple and temporally distant challenges with WT *Pb*ANKA parasites.

### Infection with IL-6 Tg-*Pb*ANKA/LISP2 SPZ confers lasting protection in a species-transcending manner

To establish whether resolved IL-6 Tg-*Pb*ANKA/LISP2 SPZ-induced parasite infection might confer protection against challenge with heterologous parasites, C57BL/6 mice infected twice with IL-6 Tg-*Pb*ANKA/LISP2 SPZ and challenged one month later with WT *Pb*ANKA SPZ and which were completely immune to reinfection, were infected 4 months later either with homologous WT *Pb*ANKA SPZ or with heterologous nonlethal *P. yoelii* 17XNL SPZ (Fig. 7A). In contrast to the control group of mice infected with *P. yoelii* 17XNL SPZ (Fig. 7B, group 3) which developed parasitemia that steadily increased to reach a peak of around 20% at day 15 post-infection, mice that were previously immunized with IL-6 Tg-*Pb*ANKA/LISP2 SPZ and challenged with *P. yoelii* 17XNL SPZ showed only a limited parasitemia (1.8%) which resolved rapidly and spontaneously until clearance by day 13 post-infection (Fig. 7B, group 2). It must be pointed out that mice challenged with the homologous WT *Pb*ANKA SPZ (Fig. 7B, group 1) did not develop any parasitemia, confirming our previous results. These data clearly demonstrate that priming with IL-6 Tg-*Pb*ANKA/LISP2 SPZ protected mice not only against homologous but also against heterologous parasite infections.

**Figure 7:**
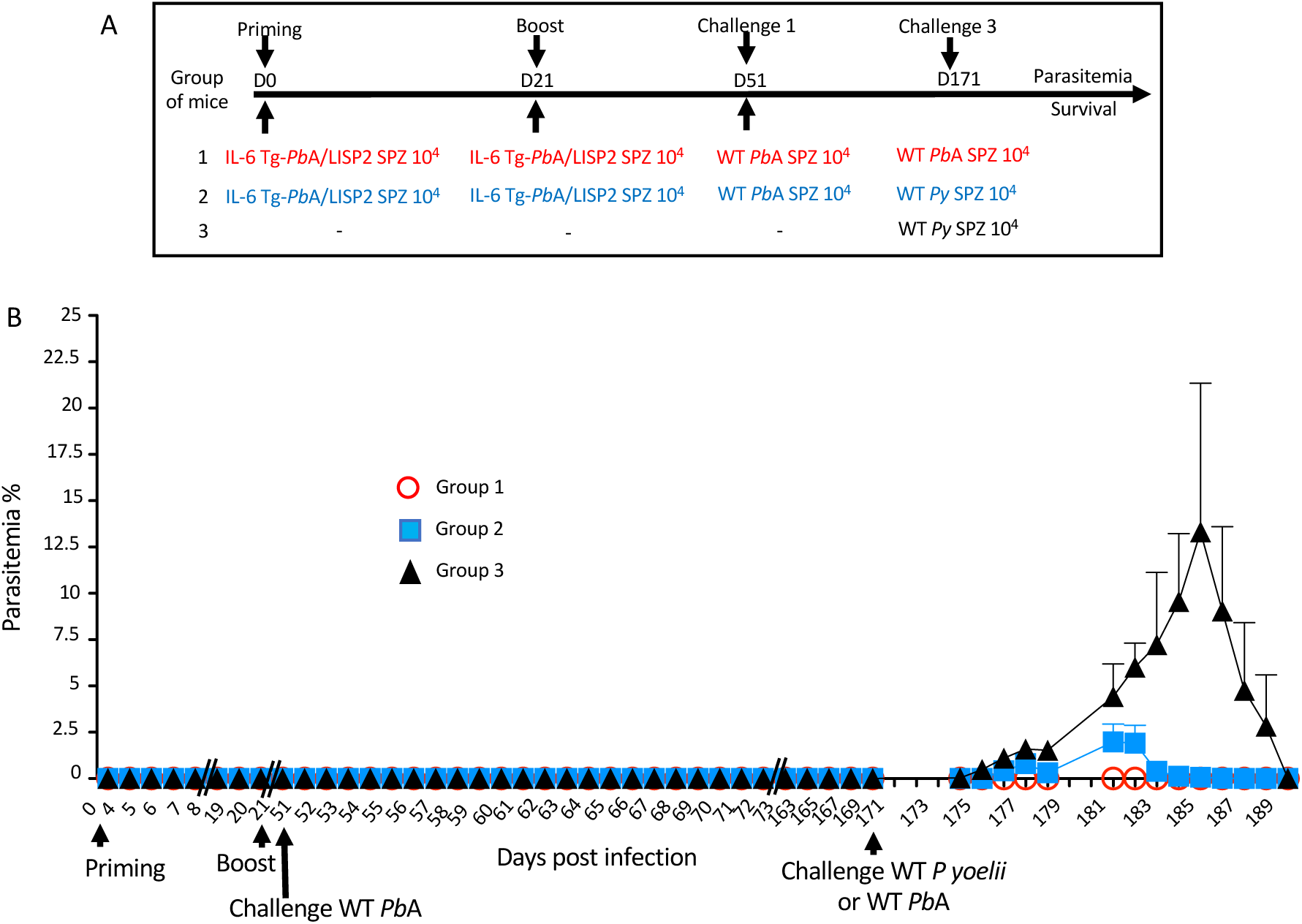
prime/boost immunization regimen with IL-6 Tg-*Pb*ANKA/LISP2 parasites confers a stable and efficient protection against challenge with heterologous *P. yoelii* 17XNL SPZ. A) Groups of C57BL/6 mice were infected twice with 10^4^ IL-6 Tg-*Pb*A/LISP2 SPZ at 3 weeks interval, and then challenged 30 days later (day 51) with 10^4^ WT *Pb*ANKA SPZ. Mice were then challenged 4 months later (day 171) either with 10^4^ WT *Pb*ANKA SPZ (group 1) or with 10^4^ WT *P. yoelii* 17XNL SPZ (group 2). A control group of age-matched naïve mice received only 10^4^ WT *P. yoelii* 17XNL SPZ. B) Parasite development was measured at indicated time points by flow cytometry, as all parasites were tagged with GFP. Error bars, SEM. Data are representative of two independent experiments with 5 mice per group.

### Protection conferred by IL-6 Tg-*Pb*ANKA/LISP2 SPZ is dependent on effector CD8^+^ T cells

In order to address whether the protection induced by IL-6 Tg-*Pb*ANKA/LISP2 parasites was dependent on effector CD8^+^ and/or CD4^+^ T cells, mice previously infected twice with IL-6 Tg-*Pb*ANKA/LISP2 parasites at 3-weeks interval were treated with normal mouse IgG, anti-CD8 or anti-CD4 depleting antibodies two days before and every other day during a challenge with 10^4^ WT *Pb*ANKA SPZ, performed 30 days after the boost (Fig. 8A). After the last infection with WT *Pb*ANKA SPZ, parasite growth and cell depletion efficacy were monitored daily by flow cytometry in blood samples. Efficacy of CD4^+^ and CD8^+^ T cell depletion was continuously monitored during administration of T-cell depleting antibodies (Suppl. Fig. 5). Interestingly, the measurement of parasitemia indicated a loss of protection upon treatment of mice with anti-CD8 but not in mice treated with anti-CD4 depleting antibodies (Group 2 and 1, Fig. 8B), while as a control all mice treated with control IgG antibodies showed sterile protection (Group 3, Fig. 8B). In terms of survival, in contrast to mice treated with control IgG and with anti-CD4 depleting antibodies, which all survived (Group 3, and 1, Fig. 8C), mice depleted of CD8^+^ cells died from hyperparasitemia (Group 2, Fig. 8C). Control naive mice which received WT *Pb*ANKA parasites during challenge (Group 4, Fig. 8B, C) developed parasitemia and died from cerebral malaria. These data strongly suggest that protection induced by IL-6 Tg-*Pb*ANKA/LISP2 parasites relies on CD8^+^ T cells but not CD4^+^ T cells.

**Figure 8:**
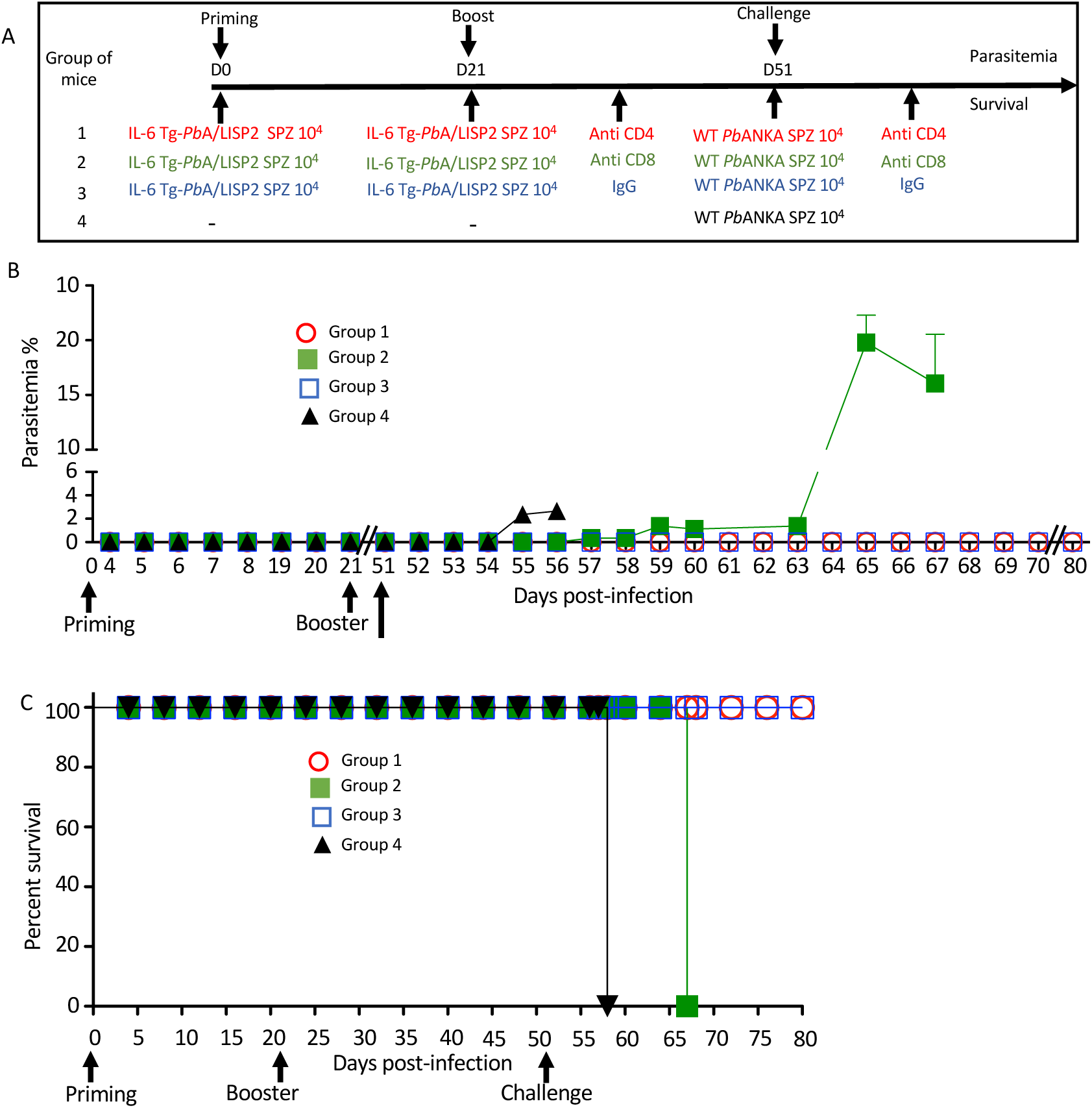
IL-6 Tg-*Pb*ANKA/LISP2 SPZ-mediated protection is dependent on CD8^+^ T cells. A) Groups of C57BL/6 mice were infected twice with 10^4^ IL-6 Tg-*Pb*A/LISP2 SPZ at 3 weeks interval, and then challenged 30 days later (day 51) with 10^4^ WT *Pb*ANKA SPZ. Two groups of mice were treated with either anti-CD4 (group 1) or anti-CD8 (group 2) depleting antibodies 2 days before and every other day after challenge. Control mice (group 3) were treated with non-specific IgG. B) Parasitemia was monitored over time by FACS analysis, as parasites were tagged with GFP. C) Survival rates were determined by Kaplan-Meier survival plots. Error bars, SEM. Data are representative of two independent experiments with 5 mice per group.

### IL-6 Tg-*Pb*ANKA/LISP2-induced protection is associated with the accumulation of memory T cells

To examine if sterile protection against malaria infection and disease induced by IL-6 Tg-*Pb*ANKA/LISP2 parasites was associated with the development of memory T cells, we analyzed the expansion and the phenotype of liver CD4 and CD8 T cells following a recall response one week after mice were inoculated twice with 10, 000 IL-6 Tg-*Pb*ANKA/LISP2 SPZ at three weeks interval (Group 1, G1, Fig. 9B) in comparison to mice infected with WT *Pb*A SPZ (Group 2, G2, Fig. 9B) and to naive mice (Group 3, G3, Fig. 9B). According to the gating strategy shown in Fig. 9A, cells were first gated on CD45^+^ CD3^+^ cells, then positive cells were gated on the expression of CD44 and CD25 markers. CD44^+^ CD25^−^ T cells were then gated on either CD4^+^ or CD8^+^ T cells. The absolute number of CD4 (panel b) and CD8 (panel d) were much higher in protected mice (G1) than in infected (G2) and in naïve (G3) mice. A higher percentage of CD8^+^ T cells (panel c) but not CD4+ T cells (panel a) was also observed in G1 protected mice. CD4^+^ and CD8^+^ T cells were further gated based on the expression of CD69 and CD62L-positive cells to analyze various subsets of memory T cells: Effector memory T cells (T_EM_): CD69^−^ CD62L^−^ CD44^+^ (panels e to h), Central memory T cells (T_CM_): CD69^−^ CD62L^+^ CD44^+^ (panels i to l), and liver resident memory T cells (T_RM_): CD69^+^ CD62L^−^CD44^+^ (panels m to p). As shown in Fig. 9B, significantly higher total number of T_EM_, T_CM_, and T_RM_ CD4 (panels f, j, n, respectively), and CD8 (panels h, l, p, respectively) were observed in protected G1 mice as compared to WT *Pb*A G2 infected mice and to naïve G3 mice. These results are consistent with the efficient and long-lasting immune memory elicited by IL-6 Tg-*Pb*ANKA/LISP2 SPZ.

**Figure 9:**
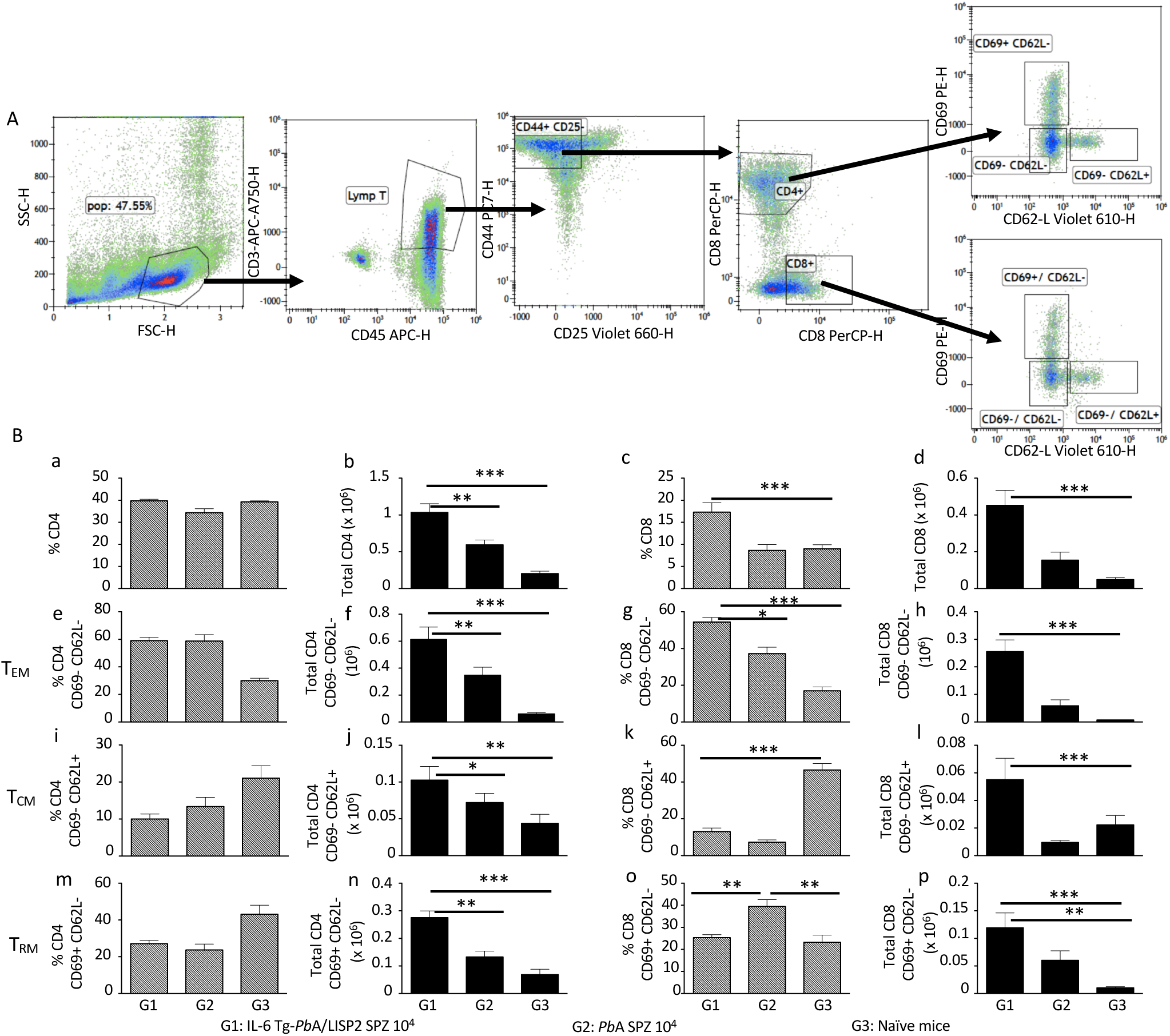
IL-6 Tg-*Pb*ANKA/LISP2-induced protection is associated with the accumulation of memory T cells. Livers were harvested one week after 6-week-old female C57BL/6 mice were infected twice with 10^4^ IL-6 Tg-*Pb*A/LISP2 SPZ at 3 weeks interval (G1) or one week after mice were infected with WT *Pb*A SPZ (G2) or from naive mice (G3) (n=6 per group). Leukocytes present in liver tissue were analyzed using the following markers: CD45, CD3, CD4, CD8, CD25, CD44, CD69, and CD62L. The gating strategy is shown in (A). Results are expressed in percentage (a, c, e, g, i, k, m, o) and in absolute number (b, d, f, h, j, l, n, p) of TEM (e, f, g, h), TCM (I, j, k, l), and T RM (m, n, o, p) positive cells (B). Results are representative for two independent experiments (Mann Whitney test; * p<0.05; **p<0.01; ***p<0.001).

## Discussion

The development of vaccines against malaria infection and disease needs improvements to overcome existing shortcomings [15]. In this study, we developed a new strategy based on the use of transgenic parasites expressing a host cytokine. Successfully devised transgenic *Pb*ANKA parasites, expressing murine IL-6 restricted to the liver stage, induced two important processes. First, the presence of the IL6 transgene impaired growth within host hepatocytes, leading to abortive development and a failure to progress toward blood stage parasites. Second, a homologous prime/boost immunization of mice with IL-6 Tg-*Pb*ANKA/LISP2 SPZ was able to protect mice against multiple and temporally distant challenges with WT *Pb*ANKA parasites and against heterologous challenge with *Plasmodium yoelii*. Collectively, our data show that a suicide strategy based on IL-6 expression by the parasite is an efficient approach not only to attenuate *Plasmodium* infection at the liver stage but also to elicit protective immunity against an infectious sporozoite challenge.

IL-6 is known to be a key cytokine involved in different arms of the immune response. It was shown to support the growth and enhancement of antibody production by B cells [16], as IL-6-deficient mice are impaired in their IgG production upon immunization [17]. On the other hand, IL-6 is critical in regulating CD4 T cell differentiation by promoting IL-4 production during T cell activation [18] and IL-21 production, an essential effector cytokine produced by Tfh cells [19]. Moreover, IL-6 in combination with IL-7 signaling promotes CD8 memory T cell generation after vaccination [20]. By enhancing locally two main features of the IL-6 cytokine, namely its pro-inflammatory and immune-stimulatory properties at the very moment when *Plasmodium* SPZ invade hepatocytes, we provide proof-of-principle that this strategy could be considered for future human vaccine strategies.

Our laboratory has gathered during the last decade experimental evidence, using various murine models for malaria, that the IL-6 response is critical in controlling the parasite growth by generating an effective anti-parasite immune response [8, 9, 7, 10]. We decided to focus our study on the IL-6 cytokine for multiple reasons. First, during inflammation, IL-6 was found to be constitutively stored by murine neutrophils [8, 21] and its secretion upon TLR activation is the major inducer of the hepatic acute phase proteins [22]. In a more recent work, we used mice deficient in the *multidrug resistance-2* gene (*Mdr2*^−/−^), which encodes the canalicular phospholipid flippase leading to a complete absence of phospholipids from bile, which were found to spontaneously develop liver injury with a typical inflammatory profile. In this model, we demonstrated that the intra-hepatocyte parasite development was impaired via an IL-6-dependent mechanism [10] and the abortive infection resulted in a long-lasting immunity in *Mdr2*^−/−^ mice against infectious SPZ. This IL-6-driven potent immune response was observed in the *Mdr*2^−/-^ mouse model we reported recently where not only total CD8^+^ and CD4^+^ cells but also CD8^+^ and CD4^+^ tissue resident memory T cells were present at a significantly higher levels in SPZ challenged *Mdr2*^−/-^ mice [10].

The use of IL-6 transgenic parasites for vaccine purposes must meet safety requirements. Accordingly, while developing this concept, the caveat that we imposed to ourselves was to avoid expression of the IL-6 cytokine systemically, *i.e* during the blood stage. Second and as *Plasmodium* hepatic stage is time frame-limited, a minimal risk of IL-6 associated side effects such as tumors due to chronic IL-6 production is expected [23]. Our strategy was to express the mouse *il-6* gene under the promoter of the liver specific protein 2 (LISP2) selected for its unique profile expression which is limited to the mid-to-late liver stage, as analysis by RT-qPCR showed that *LISP2* expression peaked at 48 hpi, and gradually decreased until reaching 60 h post infection [12]. Subsequently, production of IL-6 by transgenic parasites would be limited at this window of time, not exceeding the hepatic late stage merozoites. To assess whether *IL-6* gene transfected *Pb*ANKA parasites do indeed transcribe and secrete the mouse IL-6, we demonstrated that IL-6 was present in supernatants of HepG2 cell line cultured in the presence of IL-6 Tg-*Pb*ANKA/LISP2 SPZ but not in those cultured with WT *Pb*ANKA SPZ, indicating that the IL-6 production machinery was perfectly operational in *IL-6* gene transfected *Pb*ANKA parasites. More importantly, *in vivo* analysis showed that at the peak of IL-6 transgenic parasite growth in the liver, namely at 48h post-infection, *IL-6* mRNA transcripts were detected at the same time, suggesting an *in vivo* secretion of IL-6. This *in vivo* IL-6 production correlates with the infection pattern of mice infected with IL-6 Tg-*Pb*ANKA/LISP2 SPZ, namely, a complete blockade of the parasite development at the liver stage. These data are in accordance with the elicitation of an anti-parasite immune response by the IL-6 Tg-*Pb*ANKA/LISP2 SPZ due to the readily transcribed and secreted IL-6 early during parasite development into hepatocytes. Interestingly, no reversal of the infection phenotype of the IL-6 transgenic parasites was observed upon IL-6R blockade. These data strongly suggest that the IL-6-mediated signaling events most probably occur intracellularly and therefore are not affected by the receptor blockade of infused antibodies. There are indeed evidence that IL-6 acts as an intracrine growth factor in a variety of renal cell carcinoma cell lines which express no surface IL-6 receptors [24].

With regard to other *Plasmodium* parasites engineered to produce host cytokines, only one early study demonstrated that transgenic *P. knowlesi* parasites express bioactive IFN-γ. Unfortunately, the authors did not examine further the infectious behavior and whether this transgenic parasite was protective or not against an infectious parasite challenge [25]. In contrast to our present study, none of these reports could investigate the mechanisms underlying these partial protective capabilities.

To assess whether the inoculation of mice with IL-6 Tg-*Pb*ANKA/LISP2 SPZ was able to prime the immune system to cope with a subsequent challenge with WT *Pb*ANKA SPZ, we designed several protocols where we explored various parameters including single immunization at different dose levels, and homologous prime/boost immunization regimens with various time intervals between the immunization and the challenge with WT *Pb*ANKA SPZ, which was spanning between 30 and 60 days. Our results indicate that a single dose of IL-6 Tg-*Pb*ANKA/LISP2 SPZ, did not fully protect mice against blood stage infection. We observed, however, a significantly prolonged pre-patent period. In contrast, a full anti-disease and anti-parasite immunity were achieved after a homologous prime/boost delivery of IL-6 Tg-*Pb*ANKA/LISP2 SPZ whether the challenge with WT parasites occurred at 30 days or even at 60 days after the booster dose of IL-6 Tg-*Pb*ANKA/LISP2 SPZ. Furthermore, this protection was persistent and was maintained even after multiple challenges with WT parasites. This is indicative of a long-lasting immune memory elicited by IL-6 transgenic parasites. Protection was also observed in mice that were challenged with *P. yoelii* sporozoites, indicating that immunization with IL6-expressing parasites induces species-transcending immunity. We noticed that while induction of sterile immunity was dose-dependent, the observed liver stage-arrest of the transgenic parasites was dose-independent. This suggests that the protective immunity not only relies on the intrahepatic developmental arrest of the parasite but also on an optimal load of parasite antigens.

The failure to mount robust CD4^+^ and CD8^+^ T cell responses during natural infection via infectious mosquito bites is believed to be related to relatively low amounts of inoculated SPZ (about 100 SPZ), ultimately leading to a very limited number of infected hepatocytes, followed by a rapid release of merozoites in the bloodstream (2 days), at least in rodents. To overcome this insufficient parasite antigen availability during the short window of liver stage development, a more optimal liver stage antigen-specific CD4^+^ and CD8^+^ T cell responses can be triggered by a more persistent liver stage-arresting genetically attenuated parasite (GAP). In support of this, efficient CD8^+^ T cell responses can be elicited by late liver stage-arresting GAPs which provide a persistent source of parasite antigens [1]. In addition to CD8^+^ T cells which are critical effectors for the control of *Plasmodium*-infected hepatocytes in various species including humans, non-human primates and rodents [26], CD4^+^ T cells that recognize parasite-derived immunogenic peptides in the context of MHC class II molecules expressed on infected hepatocytes also contribute in effector mechanisms against *Plasmodium* parasite liver stages [27]. CD4^+^ T cells were also found to participate to protection against *Plasmodium* infection by inducing antibody production and macrophage activation [28, 29]. As an alternative strategy, we generated a liver stage-arresting IL-6 transgenic *Plasmodium* parasites which not only blocks blood stage infection but also conferred a long-lasting protection against virulent challenge with WT parasites. The requirement of only CD8 T cells in the protection induced by IL-6 transgenic parasites was demonstrated by the loss of control of parasite development into blood stages in mice treated with anti-CD8 but not in those treated with anti-CD4 depleting antibodies.

The defect in parasite development in IL-6 transgenic SPZ-infected mice which are completely refractory to infection even after multiple challenges with high doses of SPZ predicts that elicitation of T cells upon a successful vaccination regimen might have taken place in these mice. It is important to ask whether the abortive development of EEFs resulted in the priming of immune cells such as CD8^+^ T cells which were shown to correlate with protection against challenge with *P. berghei* in mice [30, 31] and against radiation-attenuated *P. falciparum* SPZ immunization in humans [32]. A subset of CD8^+^ T cells, termed tissue-resident memory T cells (T_rm_), has been reported to act as sentinels against invading pathogens, in particular these cells are capable of recognizing infected hepatocytes, and their depletion abrogated protection in mice [33, 34]. Our results show indeed that not only total CD8^+^ and CD4^+^ cells but also CD8^+^ and CD4^+^ T_rm_ cells were present at a significantly higher absolute number in mice infected with IL-6 transgenic SPZ than in WT *Pb*A-infected mice and in naïve mice. In search of other subsets of memory T cells, we examined the generation of CD4 and CD8 effector memory (T_em_) and central memory (T_cm_) T cells, defined on the basis of the surface expression of CD44, CD69, and CD62L after a prime/boost immunization protocol of mice infected with IL-6 transgenic SPZ. Similar to T_rm_ cells, it was striking to observe that protected mice displayed elevated absolute numbers of CD4 and CD8 T_cm_ and T_em_ in their livers even after parasites were completely cleared. These two subsets of memory T cells were previously reported in an alternative immunization protocol whereby following radiation-attenuated SPZ immunization, the overall population of intrahepatic CD8 T cells significantly increases as compared to naïve mice and protection was linked to the increase of both CD8 T_em_ and T_cm_ [35].

In conclusion, we demonstrate that transgenic *P. berghei* parasites that express and secrete the host immune modulatory factor IL-6 during liver stage development exhibit striking new features with a dual action on malaria infection outcome. First, IL-6 expression abolishes parasite growth at the pre-erythrocytic stage and second, establishes a long-lasting immune protection against a lethal parasite challenge against different rodent malaria species. Our strategy to express murine IL-6 gene under a stage-specific promoter restricts its production locally to the liver. This work opens new avenues for the development of effective live attenuated vaccines against exoerythrocytic forms of human *Plasmodium* parasites and is highly relevant to combat other pathogens that infect the liver.

## Materials and methods

### Ethics statement

All animal care and experiments described in the present study involving mice were conducted at the Institut Pasteur, approved by the “comité d’éthique en expérimentation animale” (CETEA) (Permit Number N° dap180040 issued on 2018) and performed in compliance with institutional guidelines and European regulations. A statement of compliance with the French Government’s ethical and animal experiment regulations was issued by the Ministère de l’Enseignement Supérieur et de la Recherche under the number 00218.01.

### Design of transgenic *Plasmodium berghei* parasites that express the mouse IL-6 cytokine

To generate transgenic *P. berghei* parasites expressing the murine *IL6* transgene, we used a selection-linked integration strategy [14] (Birnbaum et al., 2017). We assembled a plasmid construct, named GFP-SLI-IL6, containing two cassettes. The first cassette includes a 3’ terminal sequence of the GFP coding sequence, fused to a 2A skip peptide and the human dihydrofolate reductase (hDHFR) gene, followed by the 3’ UTR of *P. berghei calmodulin* (*CAM*) gene (Figure 1A, Suppl. Table 2A).

**Supplementary Table 2:**
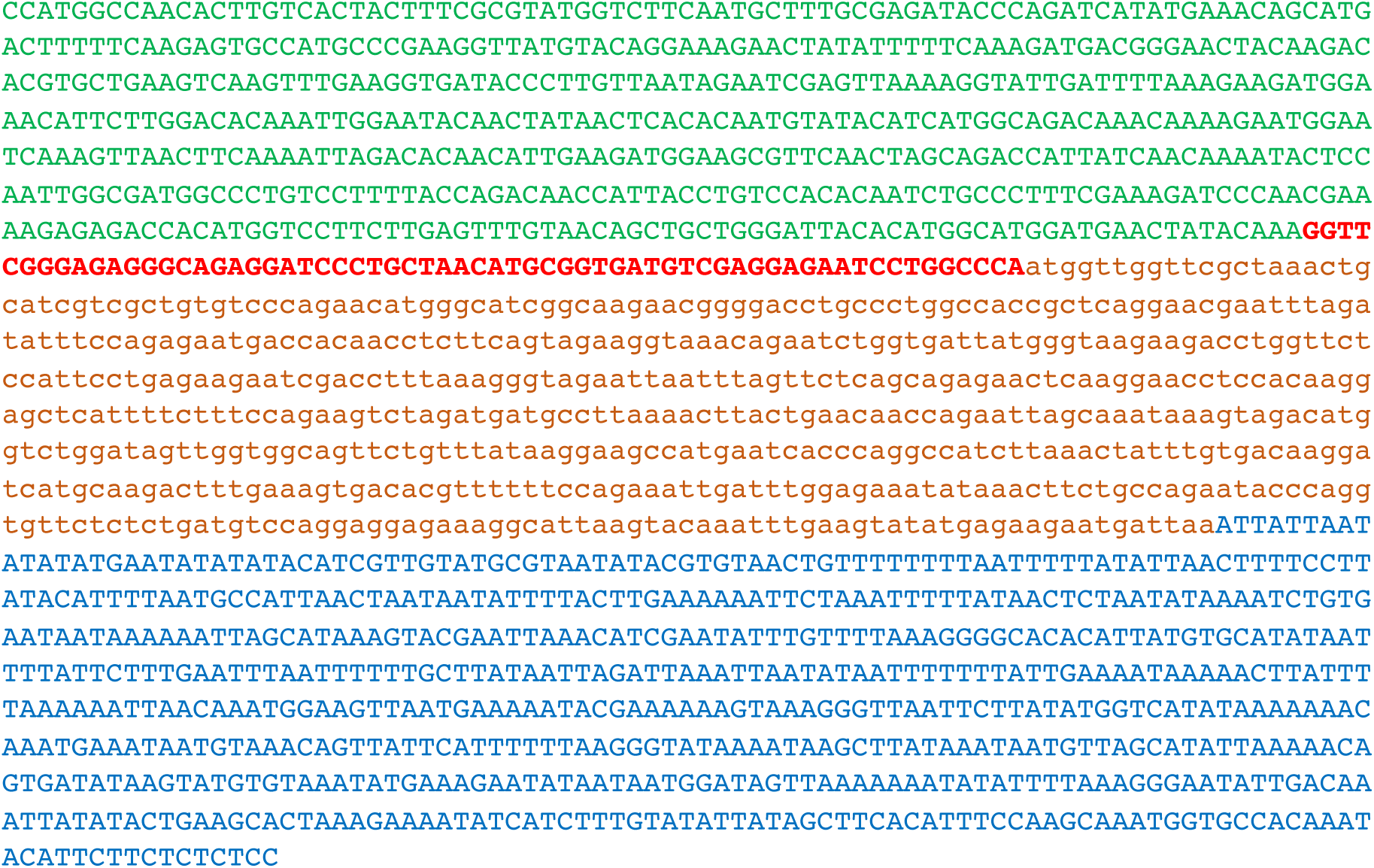
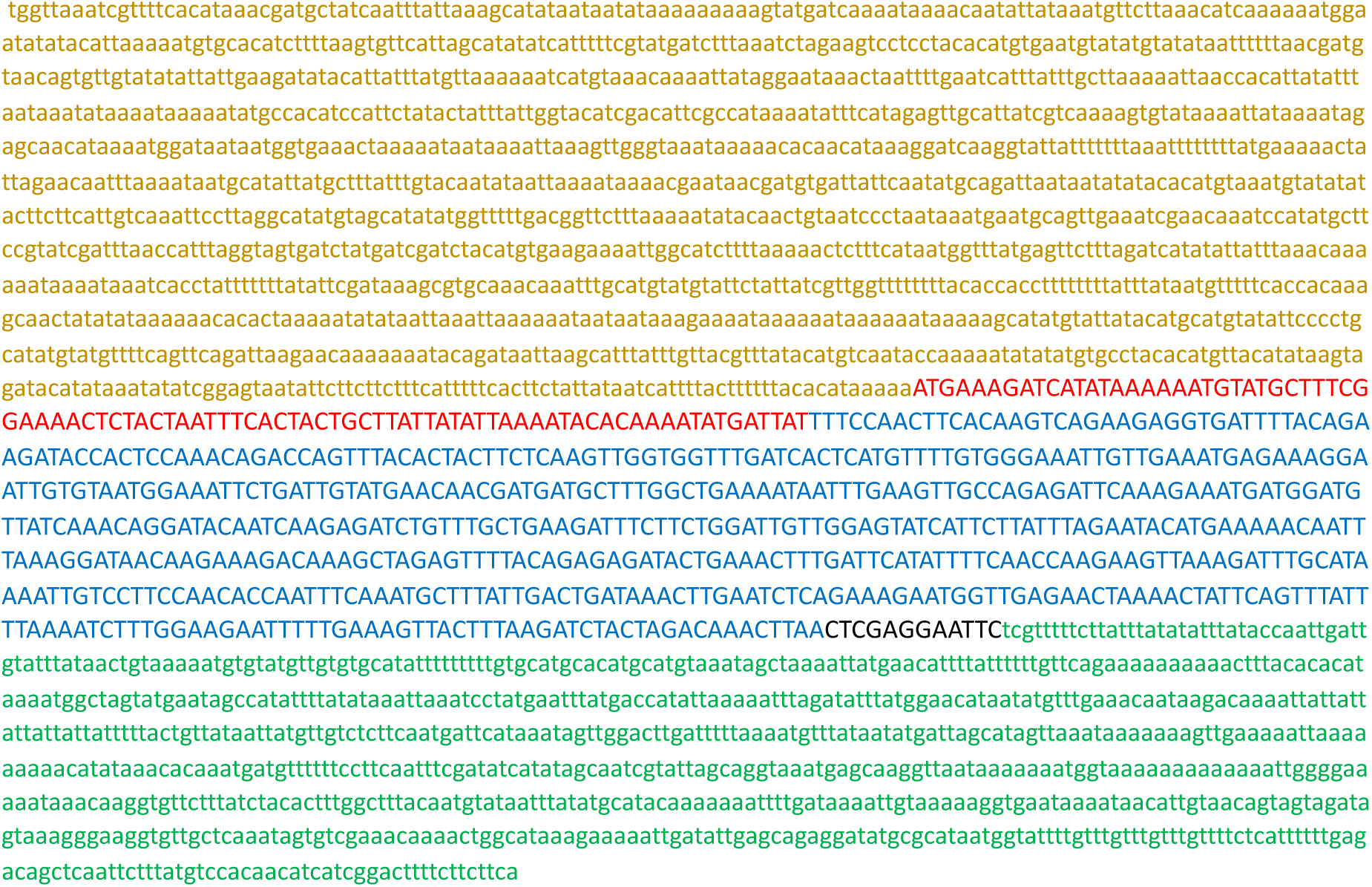
Generation of transgenic *P. berghei* parasites expressing the murine IL6 transgene using a selection-linked integration strategy which enhances locus modification through binding to the selection marker. A) Sequence of the GFP-2A-hDHFR cassette, B) Sequence of the mIL6 cassette.

**Suppl. Table 2A-first cassette** : 3’ terminal sequence of the *GFP* coding sequence, fused to a 2A skip peptide and the *human dihydrofolate reductase* gene, followed by the 3’ UTR of *P. berghei CAM*

The second cassette corresponds to a codon-optimized version of murine IL6, under control of the promoter of *P. berghei LISP2*, and followed by the 3’ UTR of *P. berghei DHFR*. To ensure IL6 secretion, the peptide signal peptide of *mIL6* was replaced by the signal peptide of *P. berghei LISP2* (Suppl. Table 2B).

**Suppl. Table 2B-second cassette** : sequence of signal peptide of *P. berghei LISP2*, codon-optimized version of murine *IL6*, under control of the promoter of *P. berghei LISP2* followed by the 3’ UTR of *P. berghei DHFR*

The final construct was verified by DNA sequencing. The linearized plasmid construct was used to transfect GFP-expressing *P. berghei* (*Pb*GFP, ANKA strain) parasites [11]. *Pb*GFP-infected erythrocytes were collected from an infected donor mice and cultured overnight to allow parasite maturation into erythrocytic merozoites. Merozoites were transfected by nucleofection with 10 μg of construct DNA, as described [36], and immediately injected into two mice. One day after transfection, mice were administered pyrimethamine in the drinking water to select for recombinant parasites. Once the parasitemia reached 2%, after one-week, infected erythrocytes were collected and DNA extracted for genotyping by PCR (Figure 1B) using primer combinations specific for the WT locus (GFP Fw + DHFRseq Rev), the 5’ integration event (GFP Fw + huDHFR Rev), the 3’ integration event (IL6 Fw + DHFRutrRev) or the non-integrated plasmid (IL6 Fw + eEF1aPromTestRev). The primer sequences are indicated in Suppl. Table 1. PCR analysis of genomic DNA from *PbIL6*/*LISP2* parasites confirmed correct integration of the constructs and the absence of residual parental parasites or non-integrated plasmid. Thus, after a single round of pyrimethamine selection following transfection, we could obtain pure populations of transgenic *PbIL6*/*LISP2* parasites. Two independent transgenic lines (line 1 and line 2) were obtained each from one of the injected mice.

### Parasites and mice infection

Seven- to eight-week-old female C57BL/6J Rj mice were purchased from Janvier Laboratories (Le Genest-Saint-Isle, France), and were maintained at the animal facility of Institut Pasteur. Mice were infected with either WT *Pb*ANKA SPZ or with IL-6 Tg-*Pb*ANKA/LISP2 SPZ or with *P yoelii* 17XNL, collected from salivary glands of infected *Anopheles stephensi*. All parasites were GFP-tagged [11]. Infections were performed via i.v. injection in the tail vein of an indicated amount of SPZ according to the experiments. In some experiments, to look whether WT and IL-6 transgenic parasite interfere with each other’s development *in vivo*, mice were co-infected with 10, 000 GFP-tagged IL-6 Tg-*Pb*ANKA/LISP2 SPZ and mCherry-tagged WT *Pb*A SPZ. Survival and parasitemia as determined by FACS using Cytoflex cytometer (Beckman Coulter Life Sciences, Villepinte, France) and the software FlowJo (FlowJo LLC, Ashland, OR, USA) were then monitored daily, beginning day 4 p.i. Symptoms associated with the experimental cerebral malaria (ECM) in mouse models include coat ruffling, a respiratory distress syndrome, a drop in body temperature, and neurological signs such as paralysis, and coma, followed by death. For ethical reasons, manifestation of signs such as coat ruffling and reduced motor skills which represent a limit point, constitutes a criterion for interrupting the experience.

### Quantification of EEF development *in vivo*

C57BL/6 mice were inoculated i.v with 150,000 WT *Pb*ANKA or IL-6 Tg-PbANKA/LISP2 SPZ and livers were harvested at 24 h and 40 h post-infection. EEF density and dimensions were determined by microscopy on freshly dissected livers using a spinning-disk confocal system (UltraView ERS, Perkin Elmer) controlled by Volocity (Perkin Elmer) and composed of 4 Diode Pumped Solid State Lasers (excitation wavelengths: 405 nm, 488 nm, 561 nm and 640 nm), a Yokogawa Confocal Scanner Unit CSU22, a Z-axis piezoelectric actuator and a Hamamatsu Orca-Flash 4.0 camera mounted on a Axiovert 200 microscope (Zeiss). Z-stacks of images spaced 3 µm apart and covering 18 to 60 µm were acquired using a EC Plan-NEOFLUAR 10x/0.3 objective (Zeiss) or a Plan-APOCHROMAT 63x/1.4 oil objective (Zeiss). Image analysis was performed with the Fiji software [37]. Z-stacks were converted to maximum intensity projections and the resulting images were segmented using the pre-implemented Triangle algorithm. EEFs were manually selected with the Magic Wand tool to determine their dimensions (area, Feret’s diameter).

### Flow cytometric analysis of liver leukocytes

Liver resident leukocytes were obtained from naïve and infected mice twice with 10, 000 GFP-expressing *Pb*ANKA or IL-6 Tg-*Pb*ANKA/LISP2 SPZ at 3 weeks interval. Briefly, mice were lethally anesthetized one weekafter the last infection with a solution of Ketamine/Xylazine and mice livers were perfused with 30 mL of PBS 1X to remove red blood cells and circulating leukocytes. After the perfusion, livers were digested *in vitro* in a collagenase D solution (0.05 %) (Roche Molecular Systems Inc., Branchburg, USA) at 37°c for 45 minutes and mixed through a 70 μm cell strainer (Falcon® Thermo Fisher Scientific Inc, Brebière, France). Cells were washed in PBS and purified by Percoll gradient: 5 mL of an 80% isotonic Percoll solution (in PBS 1X) was applied at the bottom of a 14 mL Falcon and cells were resuspended in 5 mL of 40% isotonic Percoll solution (in PBS 1X) and gently applied over the top of the tube. The cell suspension was fractioned by centrifugation at 3000 rpm during 30 minutes without brake at 4°C. After Percoll gradient centrifugation, the leukocytes were collected at the border between the two layers. Cells were washed in cold PBS, resuspended in FACS buffer containing 2 % Fetal calf serum (FCS) and 0.01 % sodium azide and counted. Obtained liver leukocytes were stained for FACS analysis according to standard protocols in a FACS buffer with the following antibodies: AF-647-labelled anti CD45 (clone 30-F11), FITC-labelled CD4 (clone H129.19), PerCP-labelled anti-CD8 (clone (53-6.7), APC-Cy 7 labeled anti-CD3 (clone 145-2C11), PeCy7-labeled anti-CD44 (clone IM7), PE-labeled anti-CD69 (clone H1 2F3), BV605 labelled anti-CD62L (clone MEL-14), and BV785-labeled anti-CD25 (clone PC61). All antibodies were purchased from BD Biosciences, San Jose, CA, USA. Cells were washed and resuspended in FACS buffer before analysis. A total of 5 × 10^5^ living or fixed cells were analysed using a Cytoflex cytometer (Beckman Coulter Life Sciences, Villepinte, France) and the software FlowJo (FlowJo LLC, Ashland, OR, USA).

### Preparation of total RNA and RT-qPCR analysis of mRNA

For kinetic analysis of *in vivo* parasite load, livers of C57BL/6J mice infected with WT *Pb*ANKA or with IL-6 Tg-*Pb*ANKA/LISP2 SPZ were surgically removed 4h, 6h, 24h, 48h, and 72h p.i., respectively. Total RNAs were extracted from the liver samples using the guanidinium-thiocyanate-phenol-chloroform method (all from Invitrogen, Waltham, MA, USA). RNA was thereafter reverse transcribed by PCR (temperature profile: 65°C for 5 min, 42°C for 50 min, and 70°C for 15 min) using 100 U of SuperScript II reverse transcriptase (Invitrogen, Waltham, MA, USA), 40 U RNase inhibitor, and 2 µM oligo (dT) 18S, HSP70, or LISP-2 rRNA primers (Eurofins MWG Operon) per sample. For detection of transcribed IL-6 mRNA by EEF, the same procedures were applied. Expression levels of diverse transcripts were analyzed by real-time RT-qPCR using Power SYBR green PCR master mix (Applied Biosystems) and various primer sets (Suppl. Table 3).

**Supplementary Table 3:**
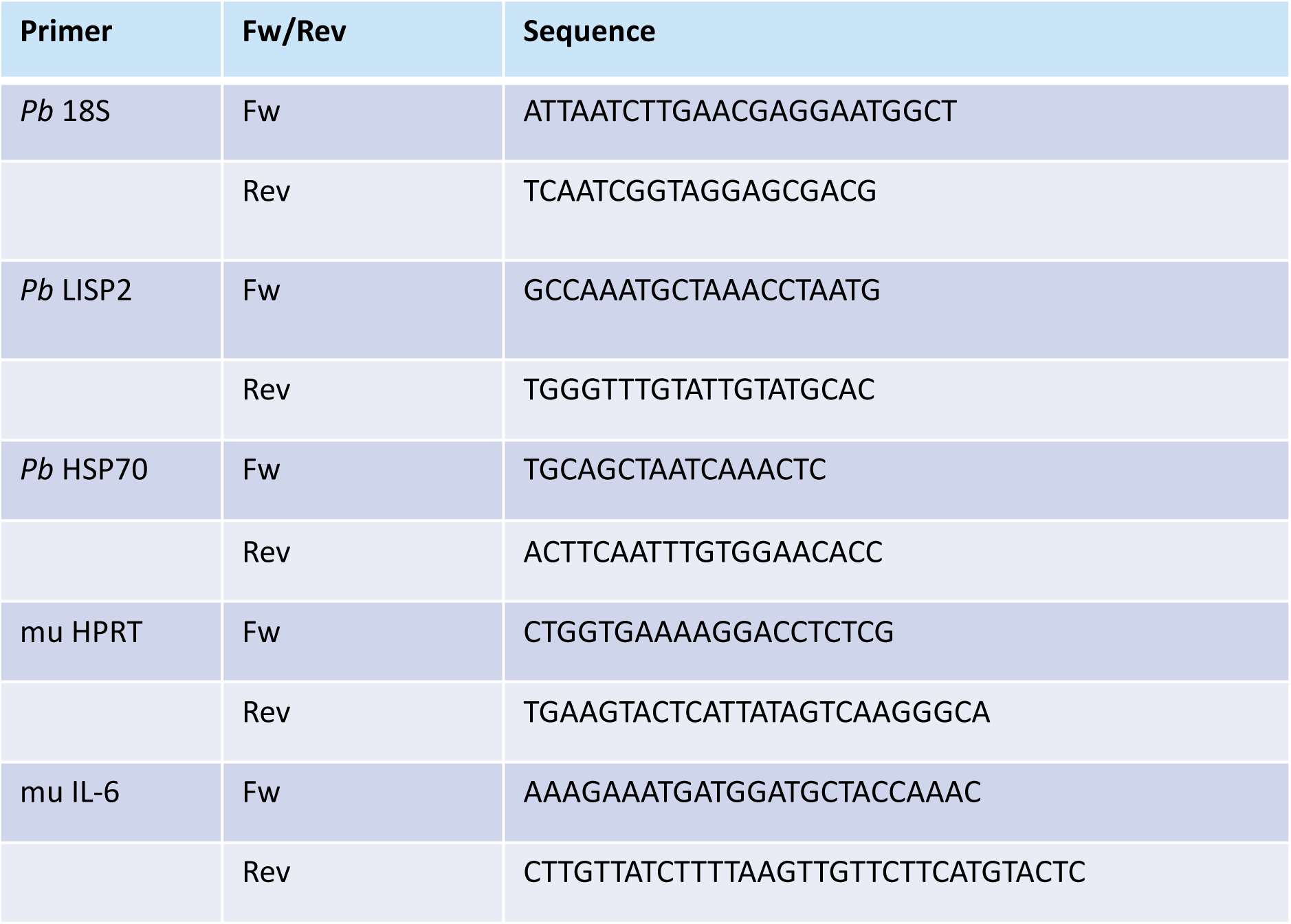
List of oligonucleotides used for RT-qPCR analyses to assess parasite load and to quantify IL-6 expression

All reactions were performed in a real-time PCR machine (temperature profile: 50°C for 2 min, 95°C for 10 min, 40 cycles of 15 s at 95°C, and 60°C for 1 min; ABI PRISM 7000 Sequence Detection System; Applied Biosystems). The relative abundance of parasite rRNA, or IL-6 mRNA in the liver and in SPZ was calculated using the ΔC_t_ method and expressed as 2−*Δ*Ct. The mouse *hypoxanthine phosphoribosyltransferase* (*HPRT*) gene was used as an internal control for the variation in input RNA amounts. A no template control was included to ensure that there was no cross-contamination during sample preparation.

### Detection of the IL-6 cytokine

HepG2 cells were infected with Tg-*Pb*ANKA/LISP2 or WT *Pb*ANKA SPZ and cultured for 48 h. Supernatants were recovered and the amounts of IL-6 were analyzed by cytokine-specific ELISA kits (BD Biosciences, Mountain View, CA).

### Treatment of mice with anti-IL-6 receptor blocking antibodies

The rat anti-mouse IL-6R antibody (clone 15A7) (BioXcell, Lebanon, NH, USA) was administered i.v at 500 μg/mouse at four consecutive times: 1 day before SPZ inoculation, and at day 0, day 1 and day 2 after SPZ inoculation. Control mice were injected with the same doses of the control isotype IgG2b (BioXcell, Lebanon, NH, USA).

### *In vivo* cell depletion

To determine if the protection induced by IL-6 Tg-*Pb*ANKA/LISP2 SPZ is dependent on effector CD4^+^ or CD8^+^ T cells, cell-specific depletion experiments were performed. C57BL/6 protected mice were injected i.p. with 20 μg of anti-CD8 clone 53-6.7 Armenian hamster IgG (eBioscience, San Diego, CA) or 100 μg of rat anti mouse CD4 clone GK1.5 (ATCC® TIB207™) 48 h before the infection with *Pb*NK65 WT followed by 6 injections administered every other day after the infection. The cell depletion was followed and confirmed every day by taking 10 μl of blood from the tip of the mouse tail and analysed by flow cytometry. Blood samples were labeled with anti-CD45-AF 647 (clone 30-F11), anti-CD8a-PE (clone 5H10) from Biolegend (San Diego, CA), and anti-CD4-FITC (clone H129-19) from BD Bioscience, Mountain View, CA.

## Statistical analysis

All data were analyzed using Prism 5.0 software (GraphPad Software, San Diego, USA). Unpaired data between two groups at a specific time point were analyzed by a Mann-Whitney test for nonparametric analysis. Kaplan-Meier survival plots were analyzed using a Mantel-Cox test. A p-value <0.05 was considered to be statistically significant. All experiments were replicated several times as indicated in the figure legends.

## Author contributions

SM: formulation of the initial concept, preparation of the original draft

SB, SB, RP, JP, P-HC, PF : performed the experiments

AS: funding acquisition and discussion

OS, SB: design and creation of the IL-6 transgenic *Plasmodium berghei* lines

OS, RA, AS: critical review and editing of the manuscript

## Abbreviations

ECM: experimental cerebral malaria
EEF: exo-erythrocytic form
GAP: genetically-attenuated parasite
*Pb*: *Plasmodium berghei*
SPZ: sporozoite
WT: wild-type

## Acknowledgements

We thank the CEPIA (Centre d’élevage, de production et d’infection des anopheles, Institut Pasteur, Paris) for providing *Anopheles* mosquitoes, and the cytometry platform (CB-UtechS facility) at the Institut Pasteur for providing technical assistance. We thank Emma Brito-Fravallo (Genetics and genomics of insect vectors Unit, Institut Pasteur) for providing technical assistance. This work has been supported by the French Parasitology consortium ParaFrap (ANR-11-LABX0024), and by a grant from Institut Pasteur to S. M.

The authors declare no competing financial interests.

## Supporting information

**Supplementary Figure 1:**
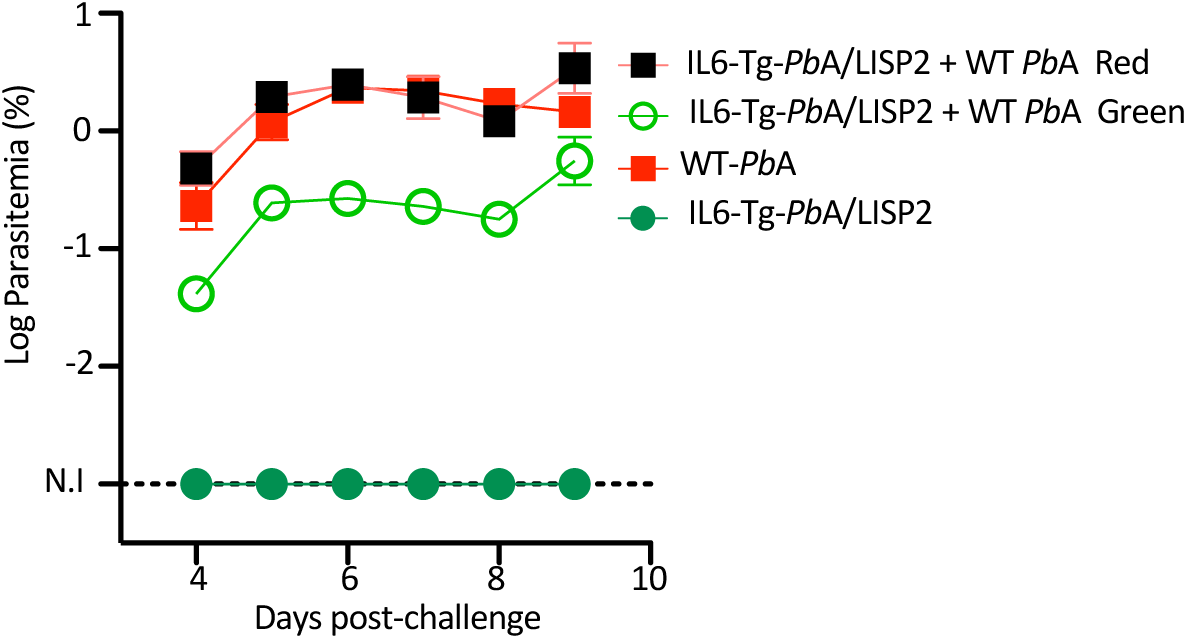
co-infection with WT parasites reversed the infection phenotype of the IL-6 recombinant parasites. C57BL/ mice were infected i.v. with a mixture of 10^4^ IL-6 Tg-*Pb*A/LISP2 SPZ and WT *Pb*ANKA SPZ. Parasitemia were measured at indicated time points by flow cytometry by counting both green and red events, as parasites were GFP tagged (IL-6 Tg-*Pb*A/LISP2) or mCherry-tagged (WT *Pb*A).

**Supplementary Fig. 2:**
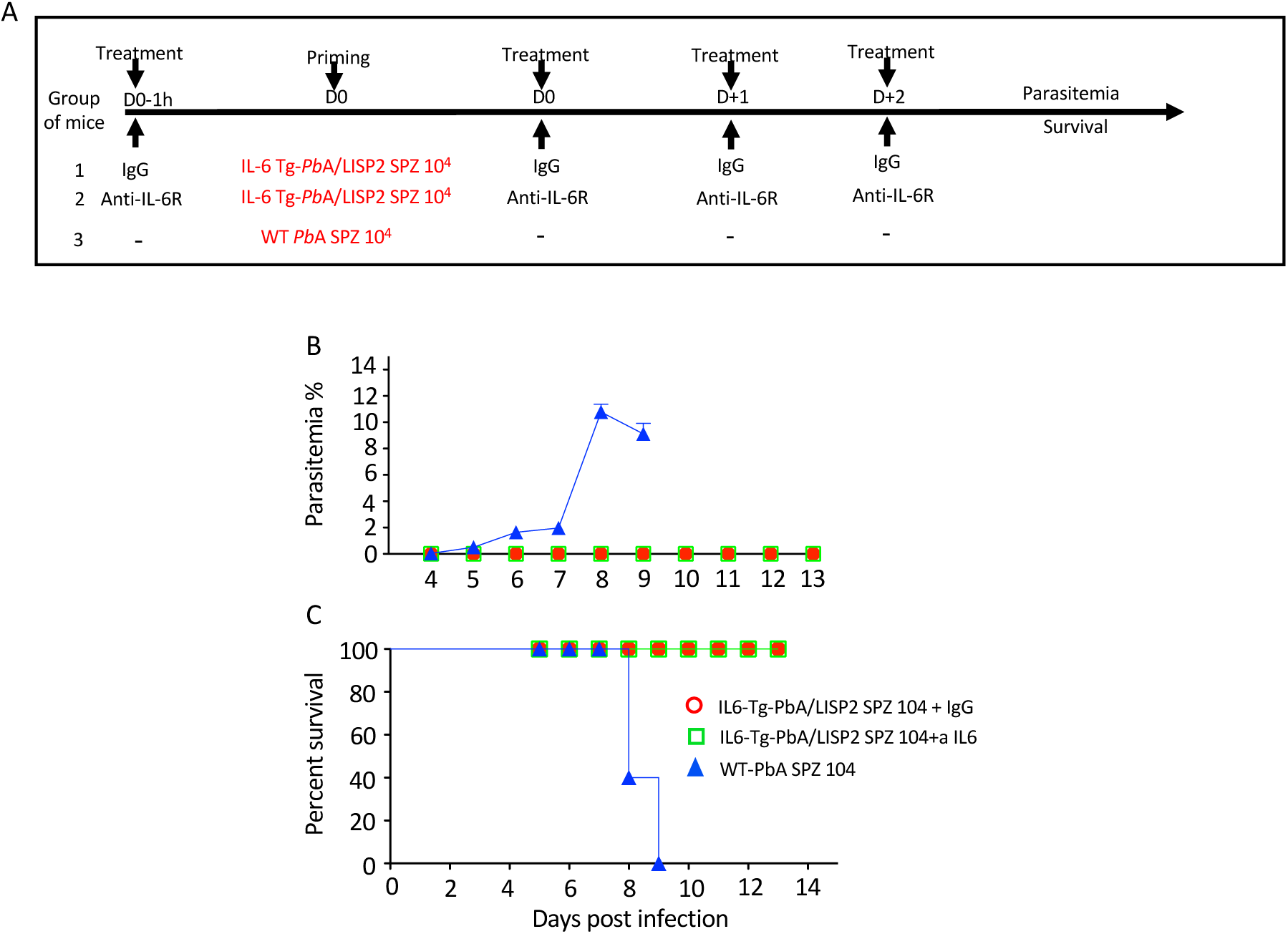
treatment with anti-IL-6R blocking antibodies does not reverse the infection phenotype of IL-6 transgenic parasites. Parasitemia (B) and survival (C) of 6-week-old female C57BL/6 mice (n=5 per group) treated i.v. with either anti-IL-6R antibody (Group 2) or with the control IgG isotype (Group 1) at 500 mg/mouse 1 day before with 10^4^ IL-6 Tg-*Pb*A/LISP2 SPZ. Mice received three additional doses of antibodies at day 0, day 1 and day 2 post-infection (A). Control mice were infected with 10^4^ WT *Pb*ANKA SPZ Group 3). Parasitemia and Kaplan-Meier survival plots (Mantel-Cox test **p<0.0061, ***p=0.001) were recorded over time. Results are from two independent experiments.

**Supplementary Figure 3:**
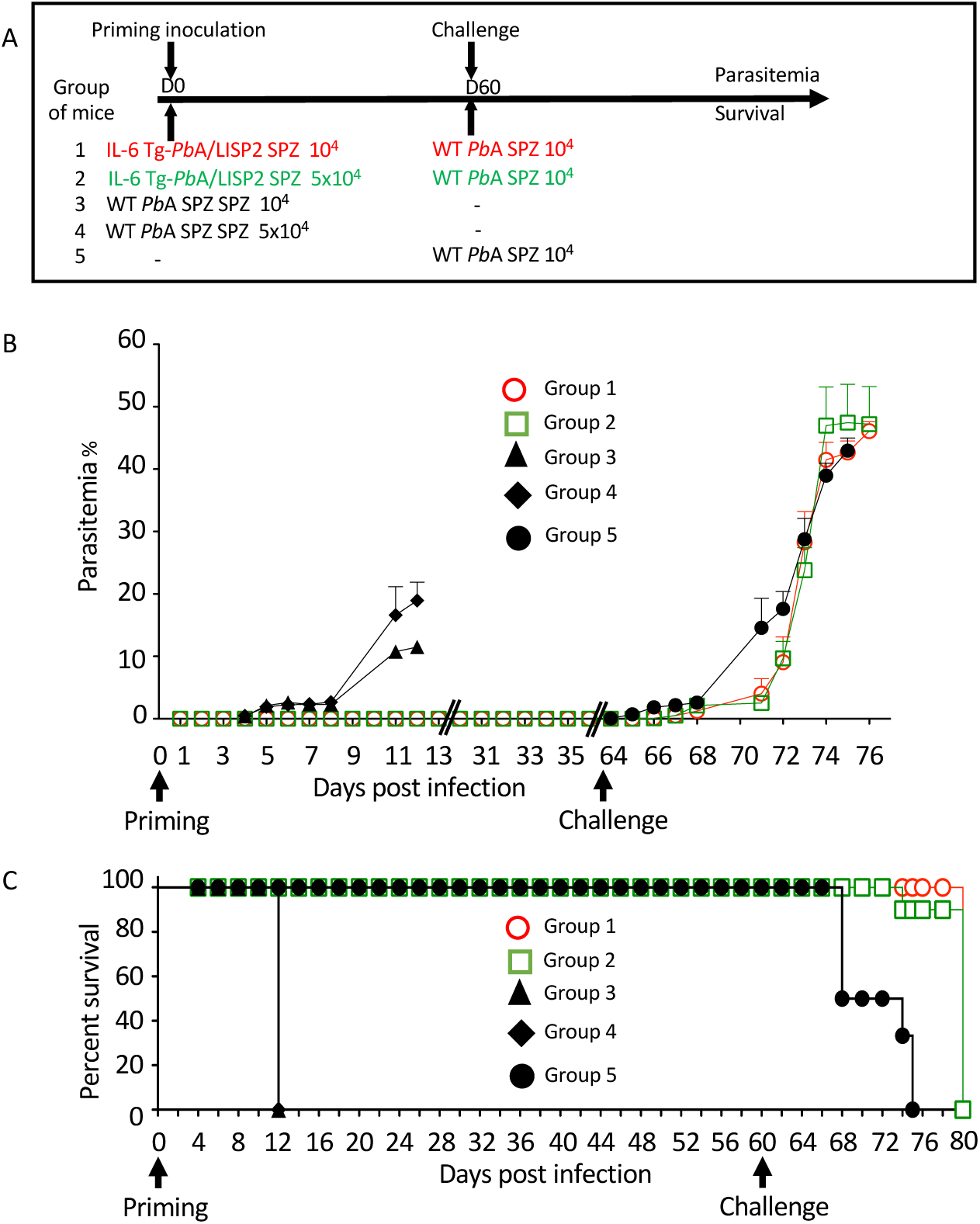
delayed challenge after priming with a single IL-6 Tg-*Pb*A/LISP2 SPZ inoculation is efficient but not fully protecting mice against lethal infection. A) Groups of C57BL/6 mice were inoculated with either 10^4^ or 5 x 10^4^ IL-6 Tg-*Pb*A/LISP2 SPZ or with the same doses of WT *Pb*ANKA SPZ. Mice were then challenged with 10^4^ *Pb*ANKA SPZ 60 days after priming. B) Parasite development was measured at indicated time points by flow cytometry, as all parasites were tagged with GFP. C) Survival rates were determined by Kaplan-Meier survival plots. Error bars, SEM. Data are representative of three independent experiments with 5 mice per group.

**Supplementary Figure 4:**
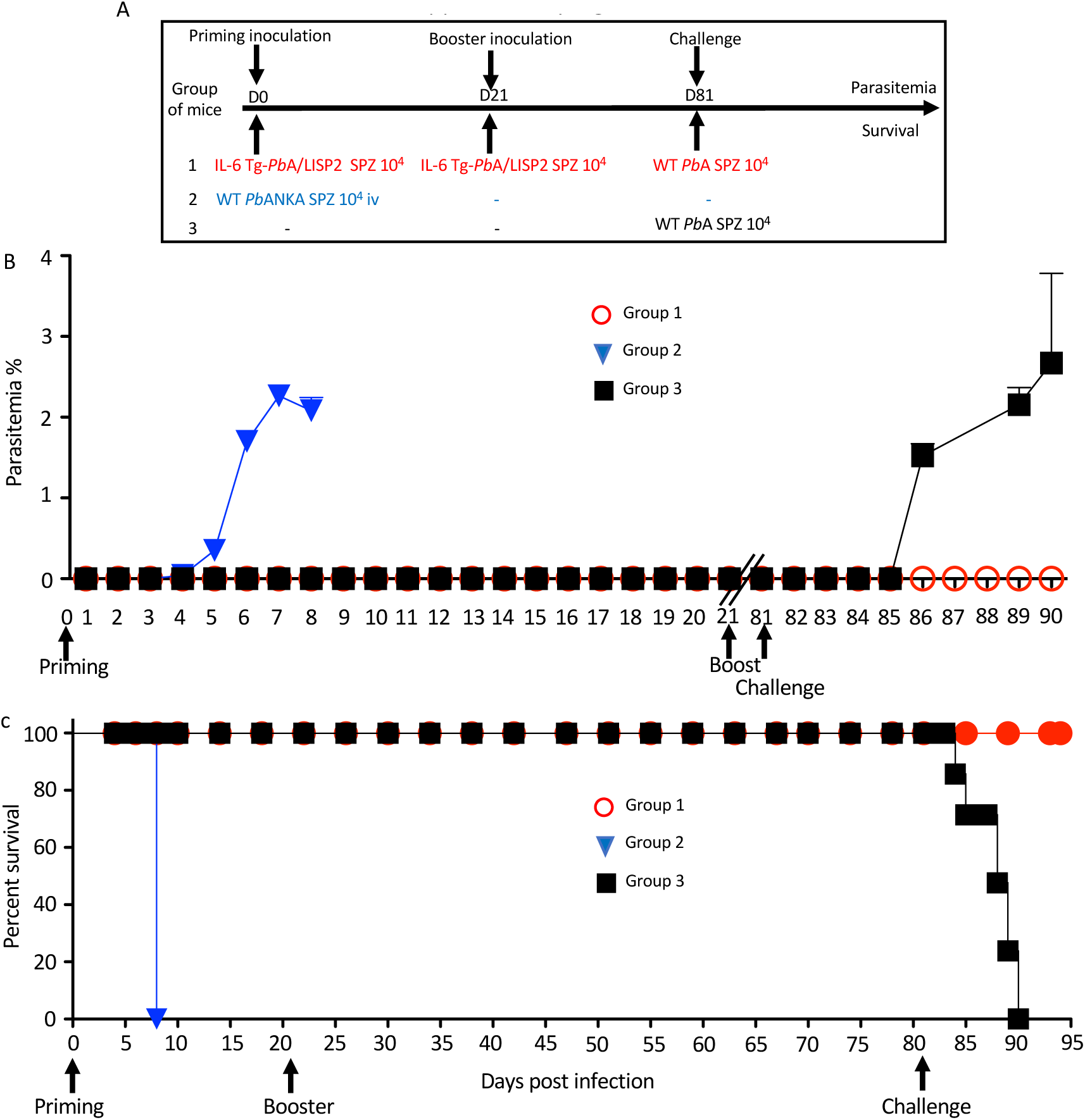
homologous prime/boost immunization regimen with IL-6 Tg-*Pb*ANKA/LISP2 parasites confers protection against temporally distant challenge with WT *Pb*ANKA parasites. A) Groups of C57BL/6 mice were infected twice with 10^4^ IL-6 Tg-*PbA*/LISP2 SPZ at 3 weeks interval or with the same dose of WT *Pb*ANKA SPZ as control. Mice were then challenged with 10^4^ *Pb*ANKA SPZ 60 days after the booster injection. B) Parasite development was measured at indicated time points by flow cytometry, as all parasites were tagged with GFP. C) Survival rates were determined by Kaplan-Meier survival plots. Error bars, SEM. Data are representative of three independent experiments with 5 mice per group.

**Supplementary Figure 5:**
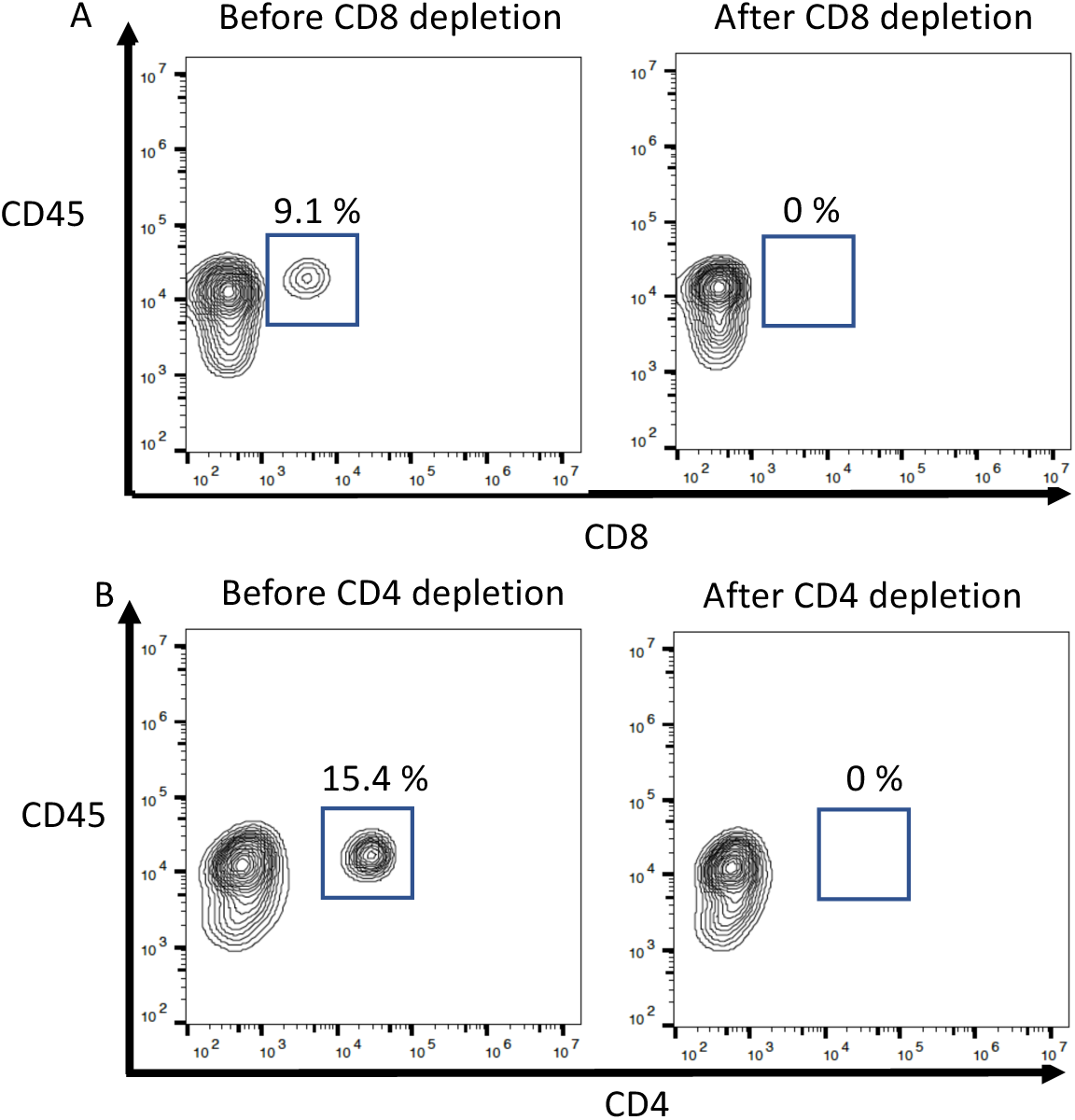
Assessment of leukocyte depletion. *In vivo* depletion of CD4^+^ or CD8^+^ T cells in immunized mice using anti-CD4 or anti-CD8 depleting antibodies was assessed by measuring daily the percentage of residual (A) CD8^+^ or (B) CD4^+^ T cells in the blood by FACS analysis. Typical analysis performed at day 2 post treatment, corresponding to the day of challenge with WT *Pb*ANKA SPZ (refer to Fig. 7), is shown in this figure. Representative data from two independent experiments with 5 mice per group are shown.

